# Chromosome-level genome assemblies of the malaria vectors *Anopheles coluzzii* and *Anopheles arabiensis*

**DOI:** 10.1101/2020.09.29.318477

**Authors:** Anton Zamyatin, Pavel Avdeyev, Jiangtao Liang, Atashi Sharma, Chujia Chen, Varvara Lukyanchikova, Nikita Alexeev, Zhijian Tu, Max A. Alekseyev, Igor V. Sharakhov

## Abstract

**Background:** *Anopheles coluzzii* and *An. arabiensis* belong to the *An. gambiae* complex and are among the major malaria vectors in Sub-Saharan Africa. However, chromosome-level reference genome assemblies are still lacking for these medically important mosquito species.

**Findings:** In this study, we produced *de novo* chromosome-level genome assemblies for *An. coluzzii* and *An. arabiensis* using the long-read Oxford Nanopore sequencing technology and the Hi-C scaffolding approach. We obtained 273.4 Mbp and 256.8 Mbp of the total assemblies for *An. coluzzii* and *An. arabiensis*, respectively. Each assembly consists of three chromosome-scale scaffolds (X, 2, 3), complete mitochondrion, and unordered contigs identified as autosomal pericentromeric DNA, X pericentromeric DNA, and Y sequences. Comparison of these assemblies with the existing assemblies for these species demonstrated that we obtained improved reference-quality genomes. The new assemblies allowed us to identify genomic coordinates for the breakpoint regions of fixed and polymorphic chromosomal inversions in *An. coluzzii* and *An. arabiensis*.

**Conclusion:** The new chromosome-level assemblies will facilitate functional and population genomic studies in *An. coluzzii* and *An. arabiensis*. The presented assembly pipeline will accelerate progress toward creating high-quality genome references for other disease vectors.

## Introduction and background

Malaria has a devastating global impact on public health and welfare, with the majority of the world’s malaria cases occurring in tropical Africa. *Anopheles* mosquitoes are exclusive vectors of malaria, with species from the *An. gambiae* complex being the deadliest African vectors. *Anopheles arabiensis* Patton, 1905 (NCBI:txid7173) and *An. coluzzii* Coetzee & Wilkerson, 2013 (NCBI:txid1518534), along with *An. gambiae* Giles, 1902 (NCBI:txid7165), are the malaria vectors of most widespread importance in Sub-Saharan Africa. *An. arabiensis* feeds and rests predominantly outdoors, replacing *An. gambiae* in some localities where there is high use of long-lasting insecticide-treated nets (LLINs) and indoor residual spraying (IRS) [1]. Females of *An. arabiensis* display opportunistic feeding behavior as they seek both human and animal blood [2]. A genomic study revealed that alleles linked to the chromosomal inversions *2Rb* and *3Ra* of *An. arabiensis* may influence choice of host for bloodfeeding [3]. Since traits relevant to vectorial capacity have genetic determinants, genomic studies are crucial for developing novel approaches to malaria control. Interspecies crosses between *An. arabiensis* and other species of the *An. gambiae* complex produce sterile males [4-6]. Fertile hybrid females allow for gene flow between species. The discovery of pervasive genomic introgression between *An. arabiensis* and *An. gambiae* or *An. coluzzii* [7, 8] opened an opportunity to investigate how traits enhancing vectorial capacity can be acquired through an interspecific genetic exchange. Hybrids of both sexes between the closely related species *An. coluzzii* and *An. gambiae* are fertile [9, 10]. These species are highly anthropophilic and endophilic; they are often sympatric but differ in geographical range [11], larval ecology [12], mating behavior, [13], and strategies for surviving the dry season [14]. Genomic analyses have the power to infer how these different adaptations are determined and maintained. A recent study described a new taxon, designated *Anopheles* TENGRELA, that genetically is most similar to *An. coluzzii* [15]. Still undiscovered and misidentified cryptic taxa could seriously confound ongoing genomic studies of *Anopheles* ecology and evolution of insecticide resistance.

The quality of genome annotation and analyses of any organism highly depends on the completeness of the assembly [16, 17]. Draft genome assemblies of species with highly-polymorphic genomes, such as mosquitoes, may have many gene annotation problems: genes can be missing entirely, have missing exons or gaps, or be split between scaffolds. As a consequence, it is difficult to estimate the total gene number or gene copy number, both of which may be linked to important phenotypic traits. Genes of particular interest with respect to vectorial capacity are especially prone to such errors since they often belong to gene families: aquaporins, ionotropic, odorant, and gustatory receptors, immunity genes, insecticide resistance genes, and reproduction gene clusters [18-25]. A genome with missing information can also cause problems for correct analyses of transcriptome, epigenome, and population genomic data. *An. gambiae*, because of its epidemiological importance, was the first disease vector that had its genome sequenced by the Sanger method (2002 and updated in 2007) [26, 27]. Since then, the AgamP4 assembly remains the standard chromosome-level genome reference for species of the *An. gambiae* complex [28-32]. Using the *An. gambiae* genome as a reference for functional annotation and population genomic analyses in other species comes at the expense of losing important information on species-specific genetic architectures in the sequencing data.

Moreover, the AgamP4 assembly has misassembled haplotype scaffolds, large gaps, incorrect orientation of some scaffolds, and unmapped sequences [26, 27]. The 16 *Anopheles* mosquito species genome project included several members of the *An. gambiae* complex [33]. Among them, the genome of *An. arabiensis* was sequenced on the Illumina platform in addition to the previously sequenced genome of the *An. coluzzii* MALI strain by Sanger [34]. Unlike the Sanger-based chromosome-level AgamP4 genome assembly, genomes of *An. arabiensis* and *An. coluzzii* MALI are represented by numerous unmapped sequencing scaffolds. Combined bioinformatics and physical mapping approaches recently produced 20 new super-scaffolded assemblies with improved contiguities for anopheline species, including *An. arabiensis* and *An. coluzzii* [35]. In the case of *An. arabiensis*, the genomic scaffolds have been assigned to and ordered and oriented on five chromosomal arms with the help of the AgamP4 reference, thus, creating the AaraD2 Illumina-based chromosome-level assembly for this species. Also, a *de novo* genome assembly from a single *An. coluzzii* Ngousso mosquito was obtained using PacBio sequencing [36]. The high-quality AcolN1 assembly for the Ngousso strain was placed into chromosome context by ordering and orienting the PacBio contigs to the AgamP4 reference. Also, 40% of the unmapped sequences in AgamP4 were assigned to the appropriate chromosomal positions [36].

Although each current sequencing or scaffolding technology alone cannot provide a telomere-to-telomere genome assembly and each method has its limitations [37], a combination of complementary approaches can lead to a chromosome-scale assembly [38, 39]. The ongoing revolution in sequencing and scaffolding methods urges researchers to undertake efforts to create new genome references that satisfy the modern standards. The Oxford Nanopore sequencing technology is a single molecule, real-time sequencing approach that utilizes biological membranes with extremely small holes (nanopores) and electrophoresis to measure the change in ionic current when a DNA or RNA molecule passes through the membrane [40]. This high-throughput technology generates exceptionally long reads and has been proven successful in genomic studies for a wide range of biological samples such as arboviruses [41], bacteria [42], plants [43], insects [44], and humans [45]. Hi-C is a groundbreaking technology that exploits *in vivo* chromatin proximity information to yield dramatically improved genome assemblies. In contrast to alternative scaffolding approaches, such as phasmid libraries or other mate-paired sequencing methods [33], the Hi-C method can produce chromosome-level genomic scaffolds [39, 46-50].

In this study, we tested the pipeline for obtaining superior-quality genome assemblies for malaria mosquitoes using the following steps: (i) high-coverage Oxford Nanopore sequencing and assembly using high-molecular-weight genomic DNA from inbred individuals, (ii) gap-filling and error-correction using Illumina sequencing data, (iii) Hi-C scaffolding of Oxford Nanopore contigs to chromosomes, and (iv) evaluation and validation of completeness and contiguity of the assemblies. We developed new reference genome assemblies for *An. coluzzii* and *An. arabiensis*, which will facilitate studies for a deeper understanding of the biology and genetics of these major African malaria vectors. The presented assembly pipeline will accelerate progress toward creating high-quality genome references for other disease vectors.

## Data Description

Here, we describe the pipeline for obtaining *de novo* chromosome-level assemblies for *An. coluzzii* MOPTI (AcolMOP1) and *An. arabiensis* DONGOLA (AaraD3). To produce the highly contiguous assemblies, we adopted and modified a strategy that was recently used for these goals in fruit flies [51, 52]. Briefly, we sequenced the genomes using the Oxford Nanopore technology, assembled contigs from long Nanopore reads, polished them with short Illumina reads, and scaffolded the contigs to the chromosome level using Hi-C proximity ligation data. Evaluation and comparison of the new assemblies with the previously released versions demonstrated substantial improvements in genome completeness and contiguity. We also provide genomic coordinates within the new references for the breakpoint regions of fixed and polymorphic inversions in *An. coluzzii* and *An. arabiensis*.

### Nanopore sequencing of *An. coluzzii* and *An. arabiensis* genomes

We performed Oxford Nanopore sequencing using genomic DNA isolated from sibling males after six generations of inbreeding to reduce heterozygosity. Our analysis and visualization of the long Nanopore reads reported 3.3 million (M) reads of the total length of 28 gigabase pairs (Gbp) and 5 M reads of the total length of 35 Gbp for *An. coluzzii and An. arabiensis*, respectively. The N50 read length was 19 kbp and 21 kbp (Additional file 1), the read median length was 3.8 kbp and 2.3 kbp (Additional file 2), the read quality was 10.3 and (Additional file 3), for *An. coluzzii and An. arabiensis*, respectively. We aligned Nanopore reads from *An. coluzzii* and *An. arabiensis* to the genome of the closely-related species *An. gambiae* (AgamP4) using minimap2 [53]. For *An. coluzzii*, the total number of aligned and unaligned reads was 3.3 M (99%) and 0.03 M (1%), respectively. In the case of *An. arabiensis*, the total number of aligned and unaligned reads was 4.5 M (89%) and 0.56 M (11%). The alignment statistics reported the 100× coverage for the *An. coluzzii* genome and the 114× coverage for the *An. arabiensis* genome (Additional file 4). Our contamination analysis with Kraken2 [54] identified 98.4% of *An. coluzzii* reads as having mosquito origin and 1.6% of reads as having bacterial origin. For *An. arabiensis*, 89.55% of the reads had mosquito origin, 5.65% of the reads had bacterial origin, and 4.8% of the reads had unknown origin. The reads of bacterial origin were filtered out. We retained the reads of unknown origin for a downstream analysis because they may represent novel mosquito sequences.

### Assembly of the *An. coluzzii* and *An. arabiensis* genomes

Genome assembly from Nanopore sequencing data is an actively developing area of research. However, no comprehensive comparisons exist of different software on diverse genomes. To assess performance of several available assemblers on *Anopheles* genomes, we ran the Nanopore reads, including wtdbg2 v1.1 [55], FLYE v2.4.1 [56], Miniasm v0.3-r179 [57], and Canu v1.8 [58]. In the case of the Canu v1.8 assembler, we obtained two assemblies: one consisting of unitigs (i.e., unambiguous reconstructions of the sequence) and the other consisting of contigs. We evaluated the contiguity of the draft assemblies using QUAST-LG [59] and estimated the genome sizes for *An. coluzzii* and *An. arabiensis* by a *k*-mer analysis of Illumina reads using Jellyfish [60]. For the *An. coluzzii* genome, the peak of the 19-mer distribution was at a depth of 54, and the genome size was estimated as 301.3 Mbp. The length of single-copy genomic regions was estimated as 204.1 Mbp (Additional file 5). For Illumina reads of *An. arabiensis*, the peak of the 19-mer distribution was at a depth of 88, and the genome size was estimated as 315.6 Mbp. The size of single-copy genome regions was estimated to be 249.4 Mbp (Additional file 5).

We obtained the following five assemblies for *An. coluzzii*:

- 1,392 contigs of a total length of 267.2 Mbp produced by Wtdbg2;
- 1,618 contigs of a total length of 279.5 Mbp produced by FLYE;
- 634 contigs of a total length of 318.0 Mbp produced by Miniasm;
- 1,073 unitigs of a total length of 344.0 Mbp produced by Canu;
- 474 contigs of atotal length of 314.2 Mbp produced by Canu.

The total lengths for Miniasm’s and Canu’s assemblies were closer to the *An. coluzzii* genome size of 301.3 Mbp, estimated from Illumina reads. According to the NG50 metric (NG50 is the length for which the collection of all contigs of that length or longer covers at least half the *An. gambiae* AgamP4 reference genome [26, 27]), the Canu contig assembly showed better contiguity (Fig. 1A, Additional file 6). However, this assembly also featured the third-largest number of misassemblies. The Wtdbg2 assembly had the smallest number of misassemblies.

**Figure 1:**
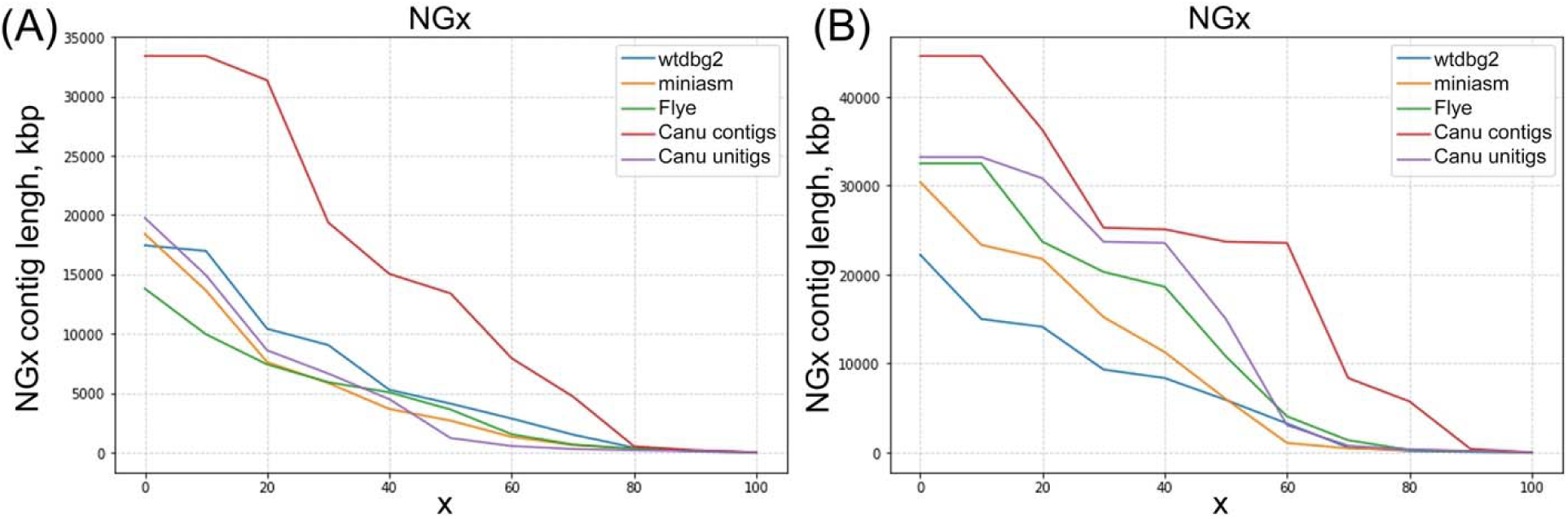
NGx curves for the Wtdbg2, Miniasm, Flye, and Canu contig and unitig assemblies. **A)** *An. coluzzii*. **B)** *An. arabiensis*.

We obtained the following five assemblies for *An. arabiensis*:

- 1,920 contigs of a total length of 298.2 Mbp produced by Wtdbg2;
- 1,280 contigs of a total length of 289.7 Mbp produced by FLYE;
- 687 contigs of a total length of 338.8 Mbp produced by Miniasm;
- 521 unitigs of a total length of 277.2 Mbp produced by Canu;
- 211 contigs of a total length of 298.5 Mbp produced by Canu.

The total length of all the assemblies except Miniasm is underestimated compared with the *An. arabiensis* genome size estimated from Illumina reads (315.6 Mbp). The *An. arabiensis* NG50 values from any assembler were substantially larger than those of *An. coluzzii* (Fig. 1B, Additional file 6). The number of misassemblies was again larger for the Canu assemblies than for other assemblies. Thus, initial assemblies of the long-read data obtained by the Canu v1.8 software alone yielded NG50 contig lengths of 13.8 Mbp for *An. coluzzii* and 23.7 Mbp for *An. arabiensis*. The total assembly sizes were 314.2 Mbp and 277.2 Mbp for *An. coluzzii* and *An. arabiensis*, respectively. For the Canu contig assemblies, the area under the NGx curve (auNG metric) was 13.57 Mbp and 21.99 Mbp for *An. coluzzii* and *An. arabiensis*, respectively, which is higher than for the other assemblies (Fig. 1, Additional file 7).

We assessed the completeness of the *An. coluzzii* and *An. arabiensis* draft assemblies with Benchmarking Universal Single-Copy Orthologs (BUSCO) v3 [61, 62]. According to the BUSCO scores for both gene datasets, Canu assemblies were the best (Additional file 8), which likely indicates that the Canu assembler has more sophisticated error-correction strategies than other assemblers have. The higher rate of core gene duplications for the Canu assemblies seems to indicate some haplotype separation, which was not observed in other assemblies. Low BUSCO scores for the Miniasm assemblies can be explained by the absence of error-correction or polishing steps in Miniasm. For example, when we polished the *An. coluzzii* Miniasm assembly with 4-round Racon [63] using Nanopore reads, the resulting assembly achieves 87.9% complete Diptera genes (Additional file 9). Based on the comparison results across all the assemblies, we decided to proceed with the Canu contig and unitig assemblies for further steps in the assembly pipeline.

### Polishing of the *An. coluzzii* and *An. arabiensis* assemblies

Initial assemblies of long reads from Oxford Nanopore Technologies are prone to frequent insertion and deletion errors, which usually are corrected by polishing. While there is no gold standard for polishing of Nanopore read assemblies, there are two commonly recommended polishing strategies. One strategy involves running several rounds of Racon [63] using raw Nanopore reads, and then running Medaka [64]. Another strategy is to run Nanopolish [65] using signal-level data additionally provided by a Nanopore sequencer. In both cases the quality of assemblies can be further improved by running Pilon [66] several times using short Illumina reads.

For polishing the *An. coluzzii* Canu contig assembly, we used Racon, Medaka, and Nanopolish, as well as Pilon for error-correction and gap-filling with Illumina reads. The FastQC quality control of the *An. coluzzii* Illumina reads reported 122.3 M reads of the total length of 22.8 Gbp and average length of 200 bp. FastQC showed that *An. coluzzii* Illumina reads have high per-base sequence quality (exceeding 32 on the Phred scale) and no adapter contamination. After each step in a polishing pipeline, we queried the resulting genome with conserved single-copy Diptera and Metazoa genes using the BUSCO test (Additional file 9). For the sake of brevity, we report only the BUSCO score for the Diptera single-copy gene set here. Single-copy genes usually cover only a small portion of a genome and it remains unclear how the polishing tools perform on the repeat-rich or non-coding regions. After Nanopolish corrected 283,935 substitutions, 1.6M insertions, and 51,104 deletions, the BUSCO score jumped from 77.6% to 93.6% of complete genes. We then ran a 4-round Racon, which dropped the BUSCO score from 93.6% to 88.6% of complete genes. Also, after Nanopolish, we ran Pilon on the Canu contig assembly several times using Illumina reads. After the first round of Pilon, we obtained 97.9% of complete genes. After three rounds of Pilon, we reached 98.5% of complete genes. Since the change was insignificant for the third round, we decided to proceed with three rounds of Pilon. We also ran Nanopolish for the second time after the first round of Pilon, but this dropped the BUSCO score to 95.9%.

For polishing the *An. arabiensis* Canu contig and unitig assemblies, we used Nanopolish and Pilon. We ran Nanopolish and 3-rounds of Pilon on the *An. arabiensis* Canu contig assembly with Illumina reads. FastQC reported 260.6 M reads for a total length 53.2 Gbp with the average length being 90 bp; further filtering out of 14% of the reads left 224.8 M of the *An. arabiensis* Illumina reads. After Nanopolish, which corrected 143,458 substitutions, 1.1 M insertions, and 40,694 deletions, the BUSCO score improved from 83% to 94.4% of complete Diptera single-copy genes (Additional file 9). After the 3-round Pilon, we obtained the Canu contig assembly with 98.5% of complete genes. We also polished the *An. arabiensis* Canu unitig assembly with Nanopolish and 3-round Pilon and obtained the BUSCO score of 98.6% of complete Diptera genes.

We remark here that the BUSCO scores for the polished Canu contig assemblies of *An. coluzzii* and *An. arabiensis* were similar to the BUSCO scores for the *An. gambiae* PEST (AgamP4) genome (Table 1). The BUSCO scores of complete Diptera genes were 98.5% for polished contig assemblies of *An. arabiensis* and *An. coluzzii*.

**Table 1:** The percentage of complete, single, duplicated, and fragmented genes computed by BUSCO from the conserved single-copy Diptera and Metazoa gene sets for the polished *An. coluzzii* and *An. arabiensis* Canu contig assemblies and the *An. gambiae* PEST genome.

| **Assemblies** | **Complete**<br>Diptera /<br>Metazoa | **Single**<br>Diptera /<br>Metazoa | **Duplicated**<br>Diptera /<br>Metazoa | **Fragmented**<br>Diptera /<br>Metazoa |
| --- | --- | --- | --- | --- |
| *An. coluzzii* Canu | 98.5% / 98.9% | 90.2% / 89.1% | 8.3 % / 9.8% | 0.7% / 0.9% |
| *An. arabiensis*<br>Canu | 98.5% / 98.9% | 93.5% / 91.8% | 5.0% / 7.1% | 0.7% / 0.2% |
| *An. gambiae* PEST | 98.3% / 98.7% | 97.7% / 96.8% | 0.6% / 1.9% | 0.8% / 0.3% |

### Hi-C scaffolding of the *An. coluzzii* and *An. arabiensis* contigs

We assembled the *An. coluzzii* and *An. arabiensis* polished Canu contigs into chromosome-level scaffolds using Hi-C Illumina reads. For both mosquito genomes, FastQC showed high per base sequence quality of Hi-C reads (exceeding 30 on the Phred scale) and detected contamination with Illumina TrueSeq adapters in 0.17% reads of *An. coluzzii* and in 1.5% of the reads for *An. arabiensis*. The contaminated reads were filtered out in both read sets, resulting in 231.9 M and 141.9 M reads for *An. coluzzii* and *An. arabiensis*, respectively. In the case of *An. coluzzii*, the Juicer tool [67] reported 3.7 M of unmapped Hi-C read pairs, 34.7 M of Hi-C read pairs mapped to inter contigs, and 97.5 M of Hi-C read pairs mapped to intra contigs. For *An. arabiensis*, we obtained 1.6 M, 9.8 M, and 66.7 M Hi-C read pairs that were unmapped, mapped to inter contigs, and mapped to intra contigs, respectively (Additional file 10).

There are several Hi-C-based scaffolding tools available: DNA Triangulation [68], LANCHESIS [69], GRAAL [70], HiRise [71], HiCAssembler [72], SALSA2 [73], and 3D-DNA [47]. Among these tools, only GRAAL, SALSA2, and 3D DNA code repositories are actively maintained. We chose 3D-DNA and SALSA2 for scaffolding of our draft assemblies because 3D-DNA allows manual correction while SALSA2 can use the assembly graphs produced by assemblers. Since SALSA2 was designed primarily for unitigs, we assessed the performance of both tools on the *An. arabiensis* unitig and contig assemblies produced by Canu. We ran SALSA2 with the corresponding assembly graph for the two *An. arabiensis* assemblies. Our experiments showed that 3D-DNA has a tendency of aggressively splitting contigs as the number of rounds grows. We, therefore, chose to run 3D-DNA with only one round instead of the default three rounds.

The QUAST-LG computation of metrics for the original and processed *An. arabiensis* assemblies demonstrated that SALSA2 performs better than 3D-DNA (Table 2). Strikingly, the assemblies processed by 3D-DNA had a higher number of scaffolds and lower NG50 than in the original assembly. At the same time, 3D-DNA corrected about 700 misassemblies in the contig assembly. While SALSA2 did not show a substantial boost in contiguity for unitig assembly, it significantly improved contig assembly at the cost of introducing a small number of misassemblies (Additional file 11).

**Table 2:** The QUAST-LG report for the NG50, number of misassemblies, and number of contigs for the *An. arabiensis* original Canu assemblies and those processed by 3D-DNA and SALSA2.

|  | **Assemblies of the *An. arabiensis* genome** |  |  |  |  |  |
| --- | --- | --- | --- | --- | --- | --- |
| **QUAST-LG metrics** | Contigs,<br>Canu | Unitigs,<br>Canu | Contigs,<br>Canu,<br>3D-DNA | Unitigs,<br>Canu,<br>3D-DNA | Contigs,<br>Canu,<br>SALSA2 | Unitigs,<br>Canu,<br>SALSA2 |
| NG50 (Mbp) | 23.7 | 23.7 | 5.1 | 5.9 | 71.4 | 23.9 |
| # misassemblies | 12,308 | 12,860 | 11,650 | 15,415 | 13,155 | 13,271 |
| # scaffolds | 211 | 521 | 246 | 870 | 161 | 366 |

We visually inspected the initial Hi-C contact heat maps produced by the scaffolding of the *An. arabiensis* assemblies (Additional file 12). The best heat map was generated by SALSA2 on the *An. arabiensis* contig assembly. For example, SALSA2 reconstructed the correct order of the scaffolds for the X chromosome except that one inversion was required to correct the orientation (upper left corner in Additional file 12a, b). On the other hand, the 3D-DNA heat maps were smoother (Additional file 12c, d) as the 3D-DNA software tends to remove repetitive sequences present in the original assemblies. We conclude that SALSA2 is better suited for scaffolding of contigs obtained from long reads. However, the results from both tools can be further improved by manual correction of scaffolds based on visual inspection of the Hi-C heat maps. Juicebox Assembly Tools (JBAT) v1.11 is the only tool currently available for manual correction of genome assemblies [74]. While 3D-DNA is designed to be loadable into JBAT for manual correction, we were unable to convert SALSA2 output to a data format that could be loaded and corrected in JBAT. Therefore, despite SALSA2 producing better scaffolding results, we decided to proceed with the 3D-DNA scaffolds obtained from the Canu contig genome assemblies for *An. arabiensis* (Additional file 12c) and *An. coluzzii* (Additional file 13).

Both species assemblies required manual correction by reordering, changing orientation, splitting contig sequences, and allocating scaffold borders. The main goal of such manual correction was to obtain chromosome-level scaffolds without assembly errors, haplotype sequences, and assembly artifacts. We also tried to minimize the number of contig splits by the manual correction.

To improve our manual correction process, we used the following additional information about contigs and scaffolds in the assemblies. All contigs were classified with PurgeHaplotigs software [75] into primary contigs, haplotigs, and assembly artifacts based on the read-depth analysis as follows. Read-depth histograms were produced for the *An. coluzzii* and *An. arabiensis* draft assemblies (Additional file 14). In each read-depth histogram, we chose three cutoffs to capture two peaks of the bimodal distribution that correspond to haploid and diploid levels of coverage. The first read-depth peak resulted from the duplicated regions and corresponded to a “haploid” level of coverage. The second read-depth peak resulted from regions that are haplotype-fused and corresponded to the “diploid” level of coverage. We also aligned contigs from each draft assembly to the *An. gambiae* PEST (AgamP4) assembly to obtain information about distribution of the contigs across the chromosomes.

Since the Hi-C signal must be stronger for adjacent sequence regions, we manually reordered and changed the orientation of contigs in each assembly to keep the Hi-C signal strong along the diagonal. We used the PurgeHaplotigs classification and the fact that haplotig sequences lead to parallel diagonal signals for moving these contigs into debris. We also moved the contigs with low Hi-C signal and the contigs classified as assembly artifacts to debris. The remaining contigs were reordered according to the Hi-C signal. After manual correction we obtained the final Hi-C contact heat maps for the chromosome-level genome assemblies of *An. coluzzii* AcolMOP1 and *An. arabiensis* AaraD3 (Fig. 2).

**Figure 2:**
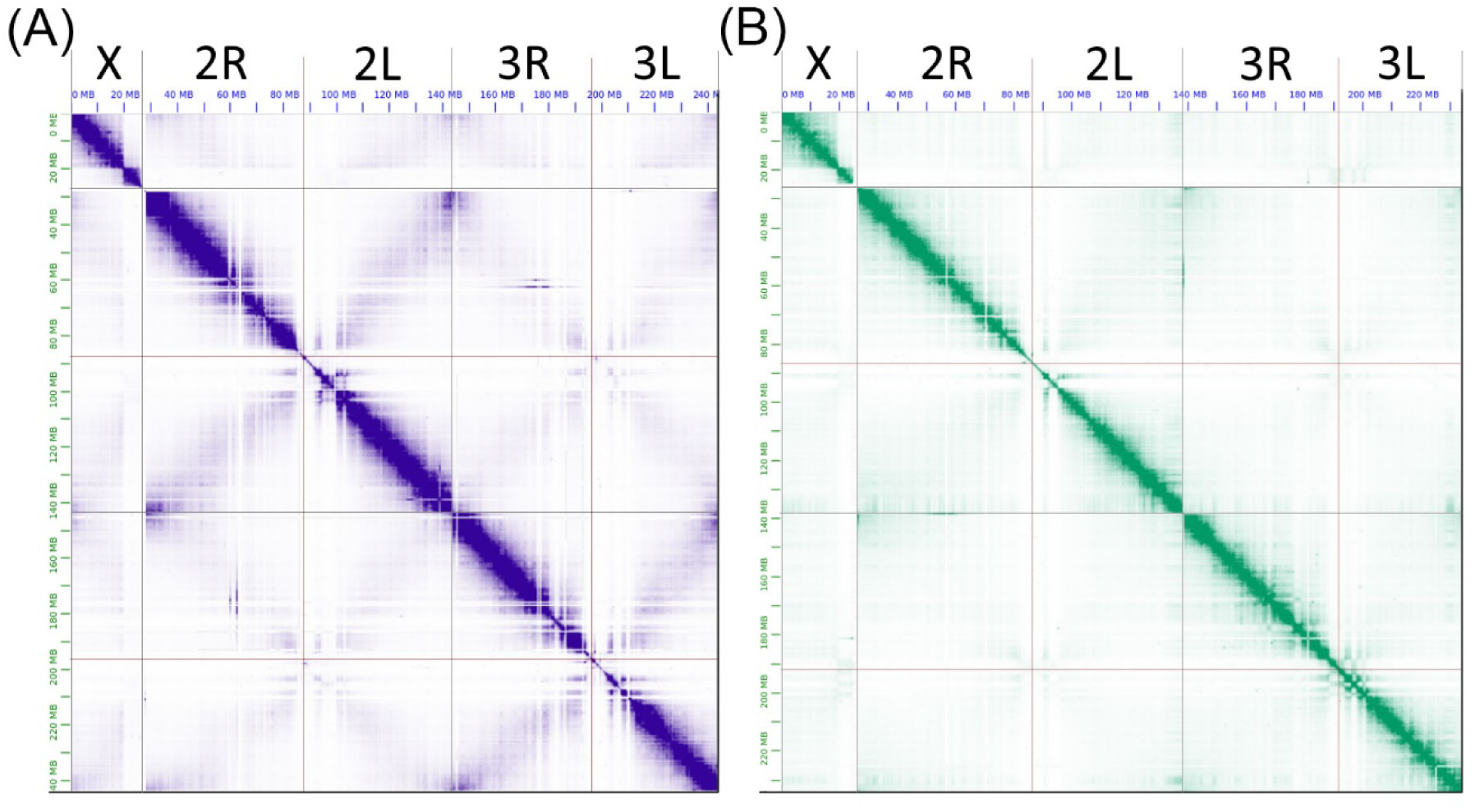
The Hi-C contact heat maps obtained after manual correction of genome assemblies. **A)** *An. coluzzii* AcolMOP1. **B)** *An. arabiensis* AaraD3. From left to right in each heat map, chromosome X, chromosome 2 (2R+2L), chromosome 3 (3R+3L). The heat maps were produced by JBAT.

Contigs in debris were further partitioned into several scaffolds. We performed chromosome quotient analysis (CQ) [76] to tentatively assign contigs in debris to the Y chromosome (CQ<0.1), X chromosome (CQ=2), and autosomes (CQ=1). We grouped contigs with CQ values less than 0.1 into the “chrY” scaffolds. In addition, we removed sequences from the chrY scaffolds if they were classified as autosomal (Ag53C, Ag93) or X chromosomal (AgX367) in the previous studies [77-79]. These results showed that the *An. coluzzii* and *An. arabiensis* assemblies contain 126 and 4 contigs from chromosome Y, respectively. We were unable to arrange contigs inside the chrY scaffolds into a correct order or orientation since the Hi-C signal was weak for these contigs.

For better understanding of the contig distribution among chromosomes, we also aligned known tandem repeats from the pericentromeric regions of chromosome X and autosomes to our assemblies. We retrieved the sequences from the debris contigs that belong to the pericentromeric region of chromosome X based on tandem repeats 18S rDNA and AgX367 and to autosomal pericentromeric regions based on satellites Ag53C and Ag93 [77-79]. Since the Hi-C signal is low for these contigs due to low complexity of the corresponding genomic regions, we were unable to determine their position inside the scaffolds forming chromosomes. Therefore, we grouped these contigs into separate “X_pericentromeric” and “Autosomal_pericentromeric” scaffolds.

As a result of the manual correction, we obtained the final assemblies for *An. coluzzii* and *An. arabiensis* genomes. They include assembled chromosomes X, 2 (2R+2L), 3 (3R+3L), and complete mitochondrion (MtDNA), as well as unordered contigs of the Y chromosome and pericentromeric sequences of autosomes and the X chromosome (including sequences from the rDNA cluster). Each species assembly consists of 7 scaffolds: chrX, chr2, chr3, chrY, X_pericentromeric, Autosomal_pericentromeric, and MtDNA. The total genome assemblies for *An. coluzzii* and *An. arabiensis* have a length of 273.4 Mbp and 256.8 Mbp, respectively (Table 3). The assembly sizes are in good agreement with the experimentally determined genome size of 260 Mbp for *An. gambiae* [80]. The resulting chromosome-level genome assemblies for *An. coluzzii* and *An. arabiensis* have a length of 242.3 Mbp and 234.6 Mbp, respectively (Table 3). By comparison, the aforementioned chromosome-level genome assemblies AgamP4 for *An. gambiae* and AaraD2 for *An. arabiensis* are 230.5 Mbp 216.6 Mbp, respectively.

**Table 3:** Genome sizes (bp) of the individual assembled chromosomes, unlocalized scaffolds, and mitochondrion for *An. coluzzii* AcolMOP1 and *An. arabiensis* AaraD3.

|  | *An. coluzzii* | *An. arabiensis* |
| --- | --- | --- |
| Chromosome X | 27656523 | 26913133 |
| Chromosome 2 (2R+2L) | 114808232<br>(60581942+54226290) | 111988354<br>(60137474+51850880) |
| Chromosome 3 (3R+3L) | 99875506 | 95710210 |
|  | (53984252+45981254) | (52890104+42820106) |
| **Total chromosome-level assembly** | **242340261** | **234611697** |
| Chromosome Y | 21271033 | 619478 |
| X_pericentromeric | 5607343 | 14883429 |
| Autosomal_pericentromeric | 4189909 | 6693365 |
| Mitochondrion | 15366 | 11309 |
| **Total genome assembly** | **273423912** | **256819278** |

### Validation and quality evaluation of the genome assemblies

We validated the resulting assemblies by aligning the AgamP4.10 gene set from the AgamP4 assembly to AaraD3 and AcolMOP1 using NCBI BLAST v2.9.0 [81]. Overall, 13,036 (99.84%) and 13,031 (99.8%) genes from the total of 13,057 *An. gambiae* genes were mapped to the *An. coluzzii* and *An. arabiensis* assemblies, respectively (Additional file 15). Moreover, 9,442 (72.31%) and 8,971 (68.71%) genes were mapped with the alignment having a zero e-value and 90%-110% of the gene length to the *An. coluzzii* and *An. arabiensis* assemblies, respectively (Table 4). The relatively smaller number of genes mapped to the X chromosome of *An. arabiensis* agrees with its higher divergence from the *An. gambiae* X chromosome [7]. The gene alignments provide supporting evidence that the obtained assemblies have the correct gene content.

**Table 4:** Statistics of the genes from the *An. gambiae* PEST assembly aligned to the *An. coluzzii* and *An. arabiensis* assemblies. In each entry (*x / y*), *x* stands for the number of genes aligned to AcolMOP1 and *y* stands for the number of genes aligned to AaraD3.

|  |  | AgamP4.10 gene set |  |  |  |  |  |
| --- | --- | --- | --- | --- | --- | --- | --- |
|  | Align<br>ed<br>from | X | 2 | 3 | Mt | Y_unplaced | UNKN |
| **Aligned to** | **total** | 1063 | 6603 | 4897 | 13 | 2 | 479 |
| **chrX** | 584 /<br>397 | 564 /<br>378 | 5 / 3 | 4 / 3 | 0 | 0 | 11 / 13 |
| **chr2** | 5055 /<br>4930 | 9 / 11 | 4862 /<br>4734 | 5 / 4 | 0 | 0 | 180 / 180 |
| **chr3** | 3785 /<br>3610 | 6 / 2 | 10 / 8 | 3566 /<br>3419 | 0 | 0 | 203 / 181 |
| **MtDNA** | 13 /<br>13 | 0 | 0 | 0 | 13 /<br>13 | 0 | 0 |
| **chrY** | 4 / 10 | 0 | 1 / 3 | 1 / 0 | 0 | 2 / 1 | 0 / 6 |
| **Autosomal_**<br>**pericentrom**<br>**eric** | 0 / 10 | 0 | 0 / 3 | 0 / 6 | 0 | 0 | 0 / 1 |
| **X_pericentr**<br>**omeric** | 1 / 1 | 0 | 0 | 1 / 0 | 0 | 0 / 1 | 0 |

We validated the structural accuracy of the AaraD3 and AcolMOP1 assemblies by comparing them with the chromosome-scale reference genome of *An. gambiae* PEST. Using D-Genies v1.2.0 [82], we generated three whole-genome pairwise alignments for the following pairs of assemblies: AcolMOP1 and AgamP4, AaraD3 and AgamP4, AcolMOP1 and AaraD3 (Fig. 3, Additional file 16). We observed that no alignment pair had inter-chromosomal rearrangements between the assemblies. Gaps in whole-genome pairwise alignments between AcolMOP1 and AgamP4 and between AaraD3 and AgamP4 indicate that all chromosomes in the new assemblies have more genomic information in the pericentromeric regions than the AgamP4 assembly. Overall, the pairwise alignments show high concordance of the AcolMOP1 and AaraD3 assemblies with the existing AgamP4 assembly.

**Figure 3:**
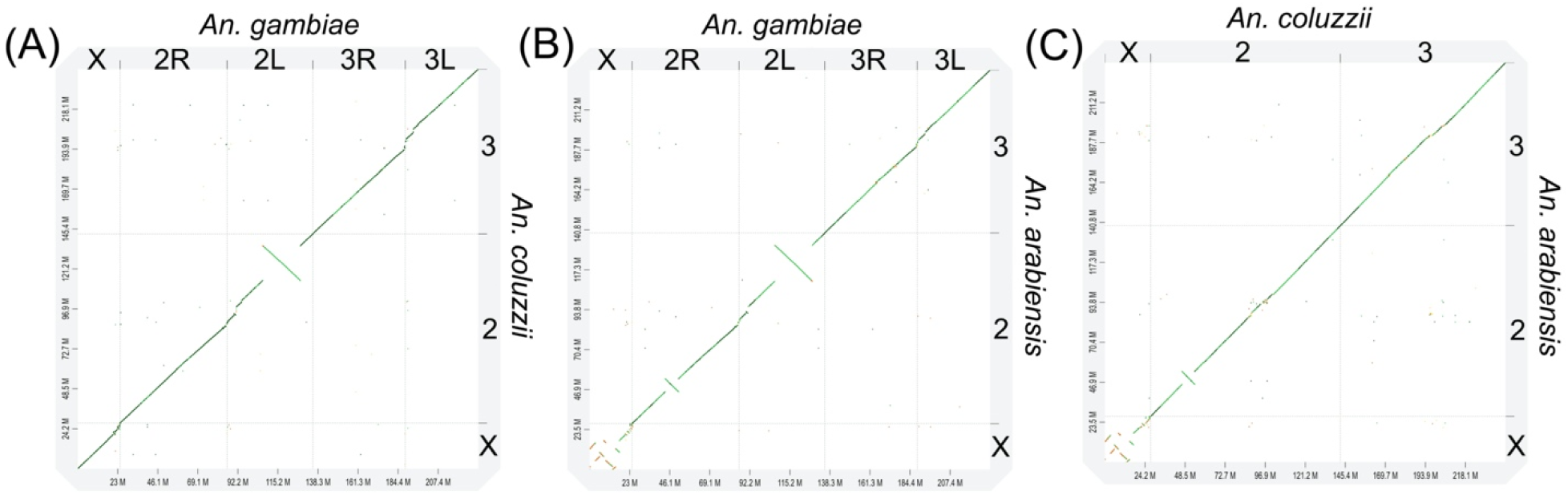
Whole-genome pairwise alignment dot-plots between chromosome-level assemblies. **A)** AcolMOP1 and AgamP4. **B)** AaraD3 and AgamP4. **C)** AcolMOP1 and AaraD3.

To assess the completeness of the final assemblies, we searched for conserved, single copy genes using BUSCO v4.1.4 with the dipteran complete gene set. The chromosomal scaffolds of the *An. coluzzii* and *An. arabiensis* final assemblies had BUSCO scores of 99.7% and 99.2%, respectively (Additional file 17). We used QUAST-LG to assess contiguity of the assemblies. The largest scaffolds are 114.8 Mbp and 112.0 Mbp and scaffold N50s are 99.9 Mbp and 95.7 Mbp for the *An. coluzzii* and *An. arabiensis* assemblies, respectively (Additional file 18).

Finally, we assessed the presence of known tandem repeats in pericentromeric regions of chromosomes 2, 3, X, and in chromosome Y of the AcolMOP1, AaraD3, AgamP4, and AcolN1 [36] assemblies. In particular, we used the presence of the putative pericentromeric tandem repeat Ag93 [78, 79] to assess the completeness of autosomal arms. We observed 27,364 and 33,460 Ag93 repeat copies in *An. coluzzii* and *An. arabiensis* assemblies, respectively, which are substantially greater than the 4,446 and 2,188 Ag93 repeat copies found in AgamP4 and AcolN1, respectively (Table 5). Moreover, while all hits in AgamP4 were located in the chromosome UNKN, 409/7003 and 1643/735 repeats were found in scaffolds corresponding to chromosomes 2/3 in the *An. coluzzii* and *An. arabiensis* assemblies, respectively (Additional file 19). This result indicates that the AaraD3 and AcolMOP1 chromosomes are more complete than the AgamP4 chromosomes. We also analyzed the presence of the Ag53C tandem repeat and its junction with the *Tsessebe III* transposable element, which are known to be located in the pericentromeric regions of the autosomes [78, 83]. The *An. coluzzii* assembly contains the highest number (646) of these repeats and the *An. arabiensis* assembly contains a comparable number of the repeats to AgamP4, while the AcolN1 assembly contains just 41 repeat copies (Table 5). It should be noted that these repeats are not located in scaffolds assembled to autosomal chromosomes and can only be found in the Autosomal_pericentromeric scaffolds of the *An. coluzzii* and *An. arabiensis* assemblies or in the chromosome UNKN of the AgamP4 assembly. Overall, the AaraD3 and AcolMOP1 assemblies contain more assembled sequences from the autosomal pericentromeric regions than the AgamP4 and AcolN1 assemblies.

**Table 5:** The number of marker sequences present in the *An. coluzzii* MOPTI (AcolMOP1), *An. arabiensis* (AaraD3), *An. gambiae* (AgamP4), and *An. coluzzii* Ngousso (AcolN1) assemblies.

| Repeat sequence | Chromosome | AcolMOP1 | AaraD3 | AgamP4 | AcolN1 |
| --- | --- | --- | --- | --- | --- |
| Ag93 | Autosomes | 27364 | 33460 | 4446 | 2188 |
| Ag53C | Autosomes | 646 | 93 | 103 | 41 |
| AgX367 | X chromosome | 845 | 1653 | 2 | 86 |
| 18S rDNA | X chromosome | 205 | 490 | 0 | 3 |
| Ag113 | X chromosome | 3146 | 0 | 765 | 97 |
| AgY477 | Y chromosome | 7685 | 0 | 4 | 0 |
| AgY53B | Y chromosome | 10569 | 0 | 387 | 0 |
| *zanzibar* | Y chromosome | 100 | 0 | 0 | 0 |

For assessing completeness of the pericentromeric regions in the X chromosome and within the Y chromosome, we used AgX367 and AgY477 repeat sequences that are known to be located in the X and Y chromosomes, respectively [77, 78]. It is important to note that AgX367 and AgY477 repeat sequences share a region of high similarity. AcolMOP1 and AaraD3 contain a much higher number of AgX367 copies than AgamP4 or AcolN1 do (Table 5). All these copies are found in the X_pericentromeric scaffold only (Additional file 19). While AcolMOP1 contains 7685 of AgY477 repeat copies in the scaffold that corresponds to the Y chromosome, they are absent in AaraD3. We also used AgY53B repeat and *zanzibar* retrotransposon to assess for the presence of Y chromosome contigs in the assemblies. Similar to AgY477, these sequences are found in AcolMOP1 but absent in AaraD3. AcolMOP1 contains the highest number of AgY53B and *zanzibar* copies among all the studied assemblies. For validating scaffolds corresponding to the X chromosome, we further used the Ag113 and 18S rDNA sequences. Remarkably, only AcolMOP1 and AgamP4 assemblies contain full-length 18S rDNA sequences. The Ag113 sequence is widely present in AcolMOP1 and AgamP4, but absent in AaraD3 (Table 5). Importanly, AcolMOP1 contains 1941 Ag113 copies in the X chromosome while all appearances of Ag113 in AgamP4 are located in the chromosome UNKN (Additional file 19). All these results indicate that the X and Y chromosomes are better assembled in AcolMOP1 than in AgamP4 or AcolN1.

We conclude that we produced chromosome-level genome assemblies for *An. coluzzii* and *An. arabiensis*. The AaraD3 and AcolMOP1 assemblies have 98.1% of conserved single-copy Diptera genes from BUSCO and contain pericentromeric heterochromatin sequences as well as sequences of the Y chromosomes and the rDNA cluster. Our assessments show that the AaraD3 and AcolMOP1 assemblies are of higher quality and more continuous than AgamP4 or AcolN1.

### Genome rearrangements in *An. coluzzii* and *An. arabiensis*

The new assemblies of *An. coluzzii* MOPTI and *An. arabiensis* DONGOLA allowed us to identify genomic coordinates of breakpoint regions and breakpoint-flanking genes of chromosomal rearrangements. The pairwise alignments of the *An. arabiensis, An. coluzzii*, and *An. gambiae* genomes were performed and visualized using D-Genies v1.2.0 [82] (Fig. 3, Additional file 17), genoPlotR [84] (Additional file 20), and SyRi [85] (Additional file 21). The karyotypes of the incipient species *An. coluzzii* and *An. gambiae* do not differ by any known fixed rearrangements. The karyotype of *An. arabiensis* is known to differ from that of *An. coluzzii* and *An. gambiae* by five fixed overlapping X-chromosome inversions (*a, b, c, d*, and *g*) and inversion *2La*. Inversions *Xag* are fixed in *An. coluzzii* and *An. gambiae* [86] and inversions *Xbcd* are fixed in *An. arabiensis* [86]. Inversion *2La* is fixed in *An. arabiensis* but polymorphic in *An. coluzzii* and *An. gambiae* [86]. Our previous studies identified the X chromosome inversion breakpoints by aligning the *An. gambiae* PEST assembly with the Illumina-based *An. arabiensis* assemblies AaraD1 and AaraD2 [7, 35]. Here, we determined genomic coordinates and breakpoint-flanking genes of the *Xag* and *Xbcd* breakpoint regions in the chromosome-level assemblies of the three species (Additional file 22). We identified breakpoint regions and breakpoint-flanking genes of inversion *2La* that is fixed in both the *An. arabiensis* DONGOLA and *An. coluzzii* MOPTI strains (Additional file 20, Additional file 21, Additional file 22). Inversion *2Rb* is polymorphic in *An. arabiensis, An. coluzzii*, and *An. gambiae* [86], and its breakpoint regions are shared among the three species indicating a single common origin of the inversion [87]. The *2Rb* inversion is fixed in the DONGOLA strain and the alignments with the *An. gambiae* and *An. coluzzii* genomes identified its breakpoints in the AaraD3 assembly (Additional file 20, Additional file 21, Additional file 22). Localization of the *2Rb* breakpoints in the context of the new reference genome will help studying of how this chromosomal inversion influences choice of host by *An. arabiensis* [3]. The genomic coordinates of breakpoint regions and breakpoint-flanking genes for inversions *2La* and *2Rb* found in our assemblies are in agreement with the previously described breakpoints for these inversions [87, 88].

Pair-wise alignments of the three chromosome-level assemblies identified new assembly-specific rearrangements, most of which are much smaller than 1 Mbp (Fig. 4, Additional file 21, Additional file 22, Additional file 23). We only considered rearrangements assembly-specific if they had the same breakpoints when aligned to both other assemblies. We found two small rearrangements in *An. coluzzii*: 3R translocation and 3L microinversion. The 3R translocation is located between genomic coordinates 37.6 Mbp and 37.7 Mbp and the 3L microinversion is located between coordinates 68.4 Mbp and 68.6 Mbp in AcolMOP1. We discovered a new 2R microinversion in *An. arabiensis* located between genomic coordinates 9.6 Mb and 9.8 Mb in AaraD3 (Additional file 21). Finally, we found five AgamP4-specific inversions on 2R (59.1–59.6 Mbp, 60.5–60.9 Mbp), on 2L (4.0–5.0 Mbp), and on 3L (0.2–0.4 Mbp, 1.2–1.9. Mbp). The identified structural variations between the assemblies may represent natural genome rearrangements or misassemblies. It is worth noticing that these micro-rearrangements are located in euchromatin of AcolMOP1 and AaraD3, while they are located in heterochromatin of AgamP4 [89]. This observation suggests that the AgamP4-specific microinversions are likely misassemblies in the PEST heterochromatin where genome assembly is notoriously difficult [26, 27].

**Figure 4:**
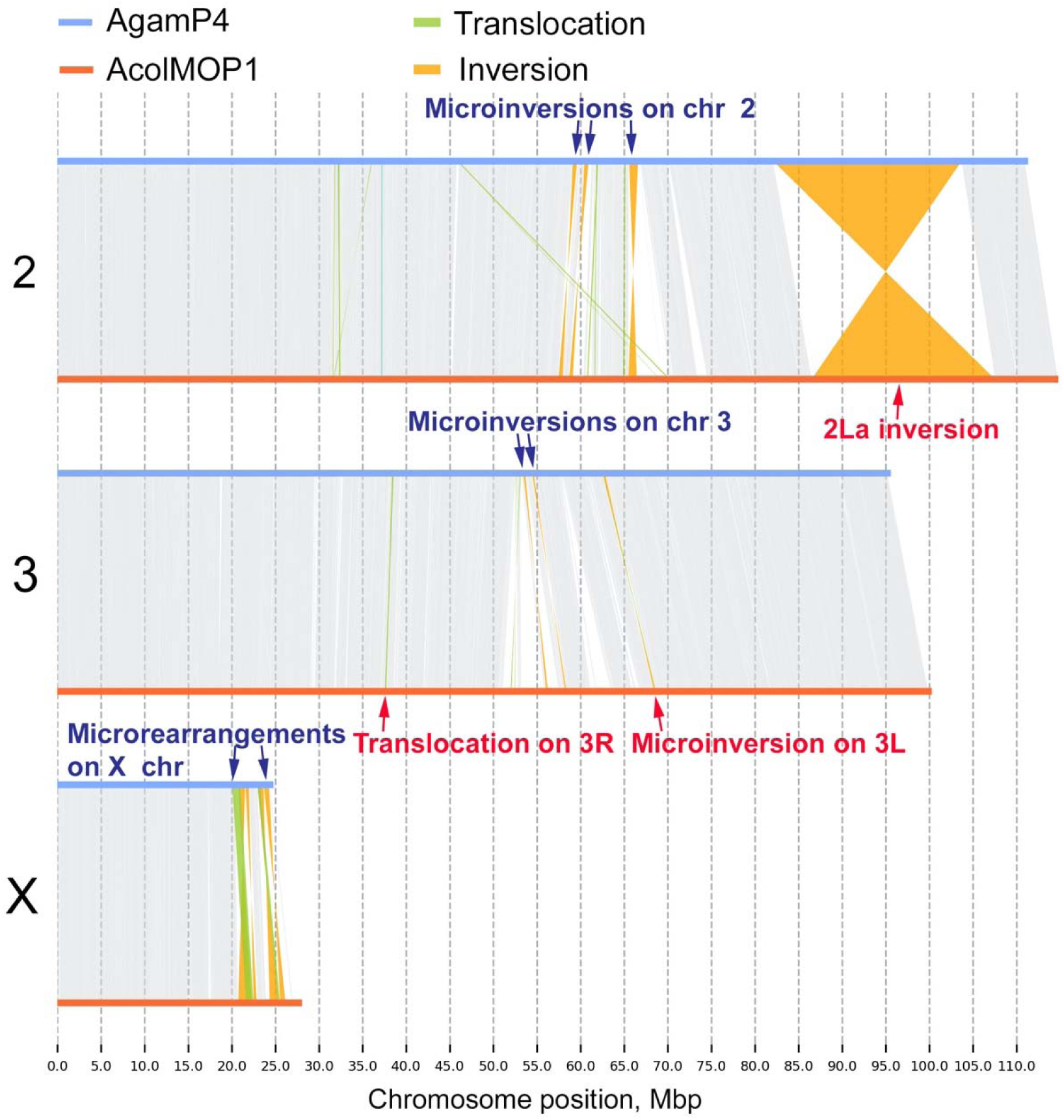
Whole-genome pairwise alignments between chromosome-level assemblies of AcolMOP1 and AgamP4. The alignment plots are produced by SyRi. The rearrangements are indicated with arrows.

To validate the new 2R microinversion in *An. arabiensis*, we aligned our AaraD3 assembly with AaraD2 [35], which is a super-scaffolded, Illumina-based AaraD1 assembly [33]. We found the new 2R microinversion in the AaraD2 assembly as well, confirming its presence the DONGOLA strain (Additional file 24). We also aligned AcolMOP1 with the reference-guided scaffolded AcolN1 assembly [36] and the super-scaffolded AcolM2 [35] assemblies. These three assemblies are made for genomes of three different *An. coluzzii* strains: MOPTI, Ngousso, and MALI, respectively. The alignments demonstrated that the new 3R translocation and 3L microinversion are MOPTI-specific (Additional file 25, Additional file 26), indicating that these rearrangements are polymorphic within *An. coluzzii*. Incidentally, we identified an inversion and two large translocations in the 2R arm (possible misassemblies) of AcolM2 in its alignment with AcolMOP1 (Additional file 26).

In addition to the rearrangements identified using these three species assemblies, we found small rearrangements in the pericentromeric heterochromatin of the X chromosome by aligning the *An. coluzzii* and *An. gambiae* genomes (Fig. 4, Additional file 27). The *An. arabiensis* genome was too divergent in this region to detect rearrangements. Breakpoint regions of two translocations and four inversions were located within genomic coordinates 20.1– 24.2 Mbp in AgamP4 and 21.6–26.2 Mbp in AcolMOP1. These rearrangements were not detected in the alignments of AcolMOP1 with either AcolN1 or AcolM2 (Additional file 27). A previous study identified three of these rearrangements in the 20–22 Mb region of the X chromosome (one translocation and two inversions) by aligning the *An. gambiae* PEST and *An. coluzzii* Ngousso genomes [36]. Since our AcolMOP1 assembly extends farther into the heterochromatin than the AcolN1 assembly does, we were able to detect two times more rearrangements in the X chromosome. These misalignments could be due to order and/or orientation errors in the PEST genome assembly. However, they may also represent novel rearrangements segregating between *An. gambiae* and *An. coluzzii*. We recently described a new type of shared cytogenetic polymorphism in the incipient species, *An. gambiae* and *An. coluzzii*—an inversion of the satDNA location in relation to the proximal gene-free X chromosome band [83]. The findings suggest that structural variations can be common in the sex-chromosome heterochromatin of mosquitoes.

## Conclusion

By combining long-read sequences generated by the Oxford Nanopore technology and long-range information produced by the Hi-C approach, we obtained high-quality reference genome assemblies for *An. arabiensis* and *An. coluzzii*. We demonstrated tremendous improvement in the completeness and contiguity of these species’ genomes. Thus, these assemblies provide a valuable resource for comparative genomics, epigenetics, functional analyses, and population studies of malaria mosquitoes. To maximize the use of the data, tools, and workflows of this study, we present a pipeline for obtaining superior-quality genome assemblies for malaria mosquitoes based on Hi-C scaffolding of Oxford Nanopore sequencing contigs (Fig. 5). The pipeline illustrates successful approaches along with other approaches that we tried but discarded in the course of its development.

**Figure 5:**
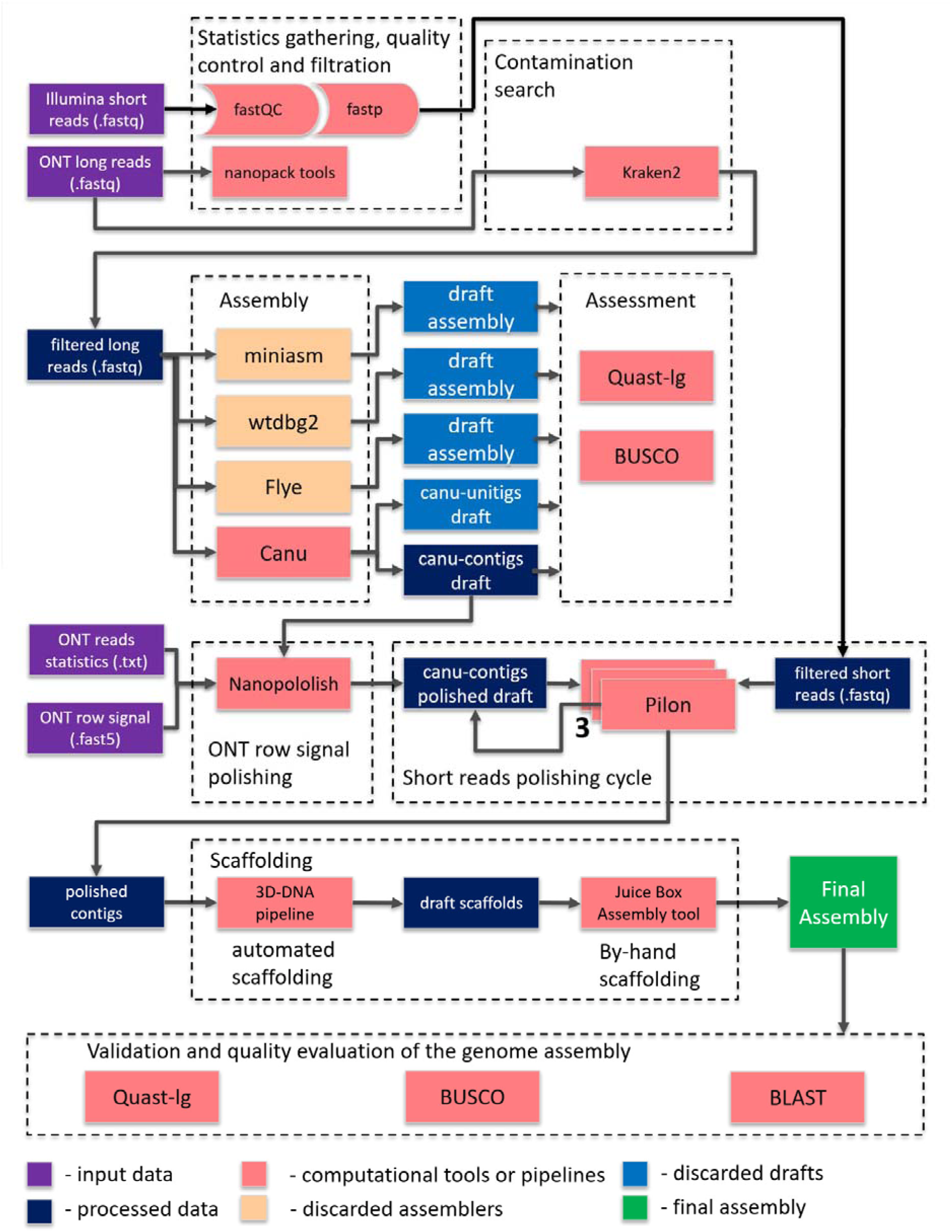
The pipeline for obtaining superior-quality genome assemblies for malaria mosquitoes based on Hi-C scaffolding of Oxford Nanopore sequencing contigs.

Sequencing of the first vector genome of *An. gambiae* revolutionized genetics and genomics research in medical entomology. *An. arabiensis* and *An. coluzzii* are major vectors of malaria, and functional characterization of their genomes will enable identification of the genomic determinants of epidemiologically important phenotypic and behavioral traits. Eventually, these efforts will lead to better malaria control.

## Materials and Methods

### Mosquito colony maintenance

The *An. arabiensis* DONGOLA (MRA-1235) and *An. coluzzii* MOPTI (MRA-763) strains were initially obtained from the Biodefense and Emerging Infections Research Resources Repository. Eggs were hatched in distilled water and incubated for 10–15 days undergoing larval and pupal developmental stages at 27 °C. After emerging from pupae, the adult males and females were maintained together in an incubator at 27 °C, 75% humidity, with a 12h cycle of light and darkness. 5–7-day-old adult females were bloodfed on defibrinated sheep blood using artificial blood feeders. Approximately 48–72 hours post bloodfeeding, egg dishes were placed for oviposition.

### Mosquito sample collection for Oxford Nanopore sequencing

A single well-blood fed female mosquito was separated from the original cage. After oviposition, F1 progeny from this single female were inbred with each other for 4 days in a 46 oz paper popcorn cup. F1 females were given bloodmeal and, after 72 hours, eggs were collected from a single F1 female. The F2 progeny were reared under normal conditions and were inbred as before. After six rounds of inbreeding using this procedure, all F7 pupal progeny from a single F6 female were sorted by sex, collected, flash frozen in liquid nitrogen, and stored at – 80°C.

### Genomic DNA isolation

Genomic DNA was isolated from 20 inbred male pupae following a modified Qiagen Genomic Tip DNA Isolation kit (Qiagen Cat No. 10243 and 19060) protocol. Briefly, pupae were homogenized using a Dremel motorized homogenizer for approximately 30 seconds at the lowest speed. Next, >300mAU (500μL of >600 mAU/ml solution) Proteinase K (Qiagen Cat No. 19131) was added to the sample and incubated at 55 °C for 3 hours. The homogenate was then transferred into a 15 mL conical tube and centrifuged at 5000 × g for 15 minutes at 4 °C to remove debris. DNA was extracted following the standard Qiagen Genomic Tip protocols. The purity, approximate size, and concentration of the DNA were tested using a nanodrop spectrophotometer, 0.5% agarose gel electrophoresis, and Qubit dsDNA assay, respectively.

### Oxford Nanopore sequencing

Approximately 1 μg of DNA was used to generate a sequencing library according to the protocol provided for the SQK-LSK109 library preparation kit from Oxford Nanopore. After the DNA repair, end prep, and adapter ligation steps, SPRIselect bead suspension (Beckman Coulter Cat No. B23318) was used to remove short fragments and free adapters. Qubit dsDNA assay was used to quantify DNA and approximately 300–400 ng of DNA library was loaded onto a MinION flow cell (MinION, RRID:SCR_017985) (BioProject: PRJNA634549, SRA: SRX8462258, SRX8462259).

### Hi-C library preparation and sequencing

Hi-C libraries for *An. arabiensis* were prepared with an Arima-HiC kit (Arima Genomics, San Diego, CA, USA) using protocols provided by the company (Document part number A160126 v00) with slight modifications. Two replicas of Hi-C libraries were prepared from 1–2-day-old virgin adults with equal proportions of each sex (one library with 20 and one library with 60 adults). After the fixation step using the *Crosslinking – Small Animal* protocol, 10% of the original pulverized mosquito tissue was taken out to perform the *Determining input Amount-Small Animal* protocol. The remaining 90% of tissue comprising at least 750 ng of DNA was used to produce proximally-ligated DNA fragments following the *Arima-HiC protocol*. The quality of proximally-ligated DNA was tested by taking 75 ng of DNA through the *Arima-QC1 Quality Control* protocol. If the Arima-Q1 value passed, the rest of the proximally-ligated DNA was used to prepare the libraries using NEBNext Ultra II DNA Library Prep Kit following the *Library Preparation* and the *Library Amplification* protocols. The libraries were sent to Novogene for sequencing. 17.8 Gb and 27.7 Gb of 2×150 bp reads were obtained for the two libraries (BioProject: PRJNA634549, SRA: SRX8462257). Three biological replicates of Hi-C data for *An. coluzzii* embryos (BioProject: PRJNA615337, SRA: SRS6448831) were obtained from the previous study [90].

### Quality control of Nanopore reads

Analysis and visualization of the long Nanopore reads were performed with the Nanostat and Nanoplot tools of the Nanopack software (de2018nanopack) [91]. Alignment of Nanopore reads to the genome of *An. gambiae* (AgamP4) was done using minimap2 (Minimap2, RRID:SCR_018550) [53]. A contamination analysis was performed with Kraken2 [54] using a custom database with addition of the *An. gambiae* [26, 27] and *An. coluzzii* Ngousso [36] genomes.

### Nanopore sequence assembly

Genome assemblies from the Nanopore sequencing data were obtained using wtdbg2 v1.1 (WTDBG, RRID:SCR_017225) [55], FLYE v2.4.1 (Flye, RRID:SCR_017016) [56], Miniasm v0.3-r179 [57], and Canu v1.8 (Canu, RRID:SCR_015880) [58]. In the case of the Canu v1.8 assembler, we obtained two assemblies: one consisting of unitigs (i.e., unambiguous reconstructions of the sequence) and the other one consisting of contigs. For wtdbg2, the polishing step using minimap2 was performed per the developers’ recommendation. The completeness and quality of the assemblies were assessed with BUSCO v2 (BUSCO, RRID:SCR_015008) [61, 62] and QUAST-LG [59]. For QUAST-LG, the *An. gambiae* (AgamP4) genome was used as a reference. *An. gambiae* is evolutionary more closely related to *An. coluzzii* (0.061 million years divergence) than to *An. arabiensis* (0.509 million years divergence) [8] and, thus, the reference-based metrics (such as NG50 and the number of misassemblies discussed below) was considered with caution. Using BUSCO, each assembly was queried for 2,799 conserved single-copy diptera genes, as well as for 978 conserved single-copy Metazoa genes. A gene recognizing model was trained for the Agustus tool [92] in the BUSCO pipeline by using the *Aedes aegypti* genome [48].

### Genome size estimation and polishing the genome assemblies

For genome size estimation and polishing assemblies obtained from Nanopore reads, Illumina short paired-end data were used. The NCBI SRX accession numbers were SRX3832577 for *An. coluzzii* and SRX084275, SRX111457, SRX200218 for *An. arabiensis*. Quality control of the Illumina reads was performed with FastQC (FastQC, RRID:SCR_014583) [93]. Based on the FastQC analysis, reads were filtered by the quality and minimum read length, and TruSeq adapters were trimmed from reads using fastp v0.20.0 (fastp, RRID:SCR_016962). The genome sizes of *An. coluzzii* and *An. arabiensis* and sizes of single-copy genomic regions were estimated by the *k*-mer analysis for *k*=19 based on the Illumina short pair-end reads using methodology described in the genome size estimation tutorial [94]. A 19-mer distribution was separated into three consecutive ranges that correspond to error sequences, single copy sequences (i.e., haploid peak), and repeat sequences. The average 19-mer single copy coverage was estimated by finding maximum in haploid peak. After that, the area under the curve was calculated for the whole distribution range except sequencing error range. For obtaining genome length, the obtained area was divided by the coverage calculated in the previous step. The length of the single copy sequences was calculated in the same manner but only for the single copy sequence range. The formulas for calculating respective lengths are the following:

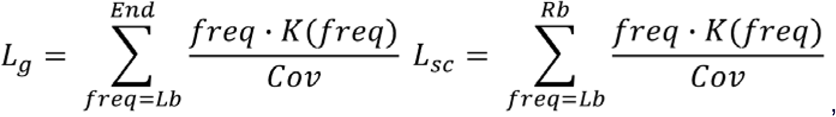

where *L*_*g*_ is a genome length, *L*_*sc*_ is a single-copy sequence length, *Cov* is an average 19-mer single copy sequence coverage, *freq* is 19-mer frequency, *K(freq)* is the number of distinct 19-mers with frequency equal *freq* (Y-axis), *End* - maximal frequency value, and *Lb, Rb are* borders of the range for haploid peak. For example, the length of single copy region for *An*.*coluzzii* genome is calculated as follows:

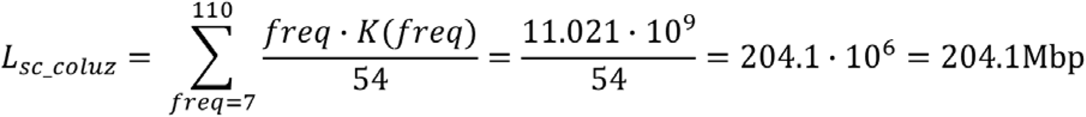

The frequency distribution of 19-mers in all high-quality short reads was computed by Jellyfish (Jellyfish, RRID:SCR_005491) [60]. Racon (Racon, RRID:SCR_017642) [63], Medaka [64], and Nanopolish (Nanopolish, RRID:SCR_016157) [65] were used to correct nucleotide substitutions, insertions, and deletions. Pilon (Pilon, RRID:SCR_014731) [66] was run several times using Illumina reads.

### Scaffolding Nanopore contigs using Hi-C data

For genome scaffolding, Hi-C Illumina short paired-end reads were used for genome scaffolding after their quality control was inspected with FastQC. BWA-MEM v0.7.17 [95] and Juicer v1.5.7 (Juicer, RRID:SCR_017226) [67] were run to assess the quality of Hi-C data with respect to the polished Canu contig assemblies of each genome. The 3D-DNA (3D-DNA, RRID:SCR_017227) [47] and SALSA2 [73] software were run to scaffold the Nanopore contigs. Metrics for the original and processed assemblies were computed by QUAST-LG [59]. Visual inspection of the Hi-C contact heat maps was performed. All contigs in the assemblies were classified with PurgeHaplotigs software [75] into primary contigs, haplotigs, and assembly artifacts based on the read-depth analysis. Alignment of contigs to the *An. gambiae* PEST (AgamP4) assembly was used for obtaining information about distribution of the contigs across the chromosomes.

### Chromosome quotient analysis

Since AgamP4 assembly does not contain chromosome Y, the chromosome quotient (CQ) analysis was performed using Illumina reads from female and male mosquito genomes to detect the presence of contigs from the Y chromosome. According to the original definition [76], for a given sequence Si, CQ(Si) = F(Si) / M(Si), where F(Si) is the number of alignments from female sequence data to Si, and M(Si) is the number of alignments from male sequence data to Si. Therefore, the CQ method allows for the differentiation of Y sequences from autosome and X sequences. CQ calculation was performed at a 1 kb window for contigs or scaffolds. If the number of male reads was below 20, the CQ value of that particular 1 kb window was not used. Contigs or scaffolds with at least 15% of the 1 kb windows showing CQ values less than 0.1 were considered as Y-derived and were grouped into a separate scaffold called “chrY.”

### Gene mapping and pairwise alignments

The *An. coluzzii* MOPTI (AcolMOP1) and *An. arabiensis* Dongola (AaraD3) assemblies were validated by comparing them with the existing assembly *An. gambiae* PEST (AgamP4), representing the most complete chromosome-level anopheline genome assembly known to date. Using NCBI BLAST v2.9.0 (NCBI BLAST, RRID:SCR_004870) [81], a set of known genes (AgamP4.10 gene set) from the AgamP4 assembly was mapped to the new *An. coluzzii* and *An. arabiensis* assemblies. The assembly of the *An. coluzzii* Ngousso strain (AcolN1) from PacBio reads was also used, where appropriate, since AcolN1 consists of contigs rather than scaffolds [36]. Whole-genome pairwise alignment between these assemblies were generated using D-Genies v1.2.0 (D-GENIES, RRID:SCR_018967) [82], SyRi [85], and genoPlotR [84]. The D-Genies v1.2.0 dot-plots and SyRi plots were generated using sequence alignments. The genoPlotR visualization was done using alignments of orthologous genes identified in the *An. coluzzii* and *An. arabiensis* assemblies by BLAST of the AgamP4.10 genes.

### Availability of supporting data and materials

Raw genomic sequence reads and genome assemblies are available in the NCBI under project accession PRJNA634549 and the *GigaScience* GigaDB [96, 97].

## Additional files

**Additional file 1**. Analysis report for Oxford Nanopore reads produced by Nanostat.

**Additional file 2**. Histograms of read length after log normalization for Nanopore reads from **(a)** *An. coluzzii* and **(b)** *An. arabiensis*.

**Additional file 3**. Plot of the average read quality for Nanopore reads obtained from **(a)** *An. coluzzii* and **(b)** *An. arabiensis*.

**Additional file 4**. Alignment depth of Nanopore reads from *An. coluzzii* (left column) and *An. arabiensis* (right column) to the *An. gambiae* (AgamP4) genome.

**Additional file 5**. Distribution of 19-mers for *An. coluzzii* (left panel) and *An. arabiensis* (right panel) computed by Jellyfish.

**Additional file 6**. Analysis report generated by QUAST-LG for Oxford Nanopore draft assemblies.

**Additional file 7**. auNG metrics for *An. coluzzii* and *An. arabiensis* assemblies.

**Additional file 8**. Evaluation of draft assembly completeness with BUSCO.

**Additional file 9**. Evaluation of assembly completeness with BUSCO after polishing steps.

**Additional file 10**. Report produced by the Juicer tool when aligning the Hi-C data on the Canu contig assemblies.

**Additional file 11**. Analysis report generated by QUAST-LG for Hi-C scaffolding of Canu contigs.

**Additional file 12**. Initial heat maps of Hi-C contact information for the *An. arabiensis* genome assemblies obtained by **(a)** SALSA 2 from the Canu contig assembly, **(b)** SALSA 2 from the Canu unitig assembly, **(c)** 3D-DNA from the Canu contig assembly, and **(d)** 3D-DNA from the Canu unitig assembly. The heat maps were produced by JBAT.

**Additional file 13**. Hi-C contact heat map for the 3D-DNA scaffolds of the *An. coluzzii* assembly before manual correction. The heat map is produced by JBAT.

**Additional file 14**. Read depth histogram obtained by Purge Haplo for the *An. coluzzii* **(a)** and *An. arabiensis* **(b)** assemblies. The cutoffs were manually selected (red arrows in the histograms): 30, 78, and 132 for *An. oluzzii* and 25, 93, and 160 for *An. arabiensis*.

**Additional file 15**. Mapping the AgamP4.10 gene set to the final *An. coluzzii* and *An. Arabiensis* assemblies.

**Additional file 16**. Whole-genome pairwise alignment dot-plots (produced by D-Genies v1.2.0) between the scaffolds corresponding to chromosomes X **(a, b, c)**, 2 **(d, e, f)**, 3 **(g, h, i)**. Alignments between the *An. gambiae* and *An. coluzzii* scaffolds, the *An. gambiae* and *An. arabiensis* scaffolds, and the *An. coluzzii* and *An. arabiensis* scaffolds are shown.

**Additional file 17**. BUSCO scores for the final assemblies.

**Additional file 18**. Analysis report generated by QUAST-LG for the final Hi-C-scaffolded Canu contigs.

**Additional file 19**. Map of the repetitive sequences for the *Anopheles* assemblies.

**Additional file 20**. Whole-genome pairwise alignments produced by genoPlotR between chromosomes of *An. arabiensis, An. coluzzii, and An. gambiae* Left panel: AgamP4 and AcolMOP1. Middle panel: AgamP4 and AaraD3. Right panel: AcolMOP1 and AaraD3. The inversion breakpoints are shown with small letters.

**Additional file 21**. Whole-genome pairwise alignments produced by SyRi of *An. arabiensis* chromosomes (query) to *An. gambiae* (reference), and *An. coluzzii* (reference) chromosomes. Left panel: AgamP4 and AaraD3. Right panel: AcolMOP1 and AaraD3.

**Additional file 22**. Genomic coordinates of breakpoint regions and breakpoint-flanking genes of chromosomal rearrangements identified by aligning the AgamP4, AcolMOP1, and AaraD3 assemblies using D-Genies v1.2.0.

**Additional file 23**. Genomic coordinates of breakpoint regions of chromosomal rearrangements identified by aligning the AgamP4, AcolMOP1, and AaraD3 assemblies using SyRi.

**Additional file 24**. Pairwise dot-plot alignment between the AaraD3 and AaraD2 (AaraD1 superscaffolded) assemblies produced by D-Genies v1.2.0. Top panel: Whole-genome pairwise alignment. Bottom panel: Collinearity in the alignment of the region with the new 2R microinversion.

**Additional file 25**. Pairwise dot-plot alignment between the AcolMOP1 and reference-guided scaffolded AcolN1 assemblies produced by D-Genies v1.2.0. Left panel: Whole-genome pairwise alignment. Middle panel: Alignment of the region with the new 3R translocation. Right panel: Alignment of the region with the new 3L microinversion.

**Additional file 26**. Pairwise dot-plot alignment between the AcolMOP1 and AcolM2 assemblies produced by D-Genies v1.2.0. Left panel: Whole-genome pairwise alignment. Middle panel: Alignment of the region with the new 3R translocation. Right panel: Alignment of the region with the new 3L microinversion.

**Additional file 27**. Pairwise dot-plot alignment between the X chromosomes produced by D-Genies v1.2.0. Left panel: AcolMOP1 and AgamP4. Middle panel: AcolMOP1 and AcolN1. Right panel: AcolMOP1 and AcolM2.

### Abbreviations

bp: base pairs;
BUSCO: Benchmarking Universal Single-Copy Orthologs;
JBAT: Juicebox Assembly Tools;
SRA: Sequence Read Archive;
NCBI: National Center for Biotechnology Information;
M: million;
Gbp: gigabase pairs;
Mbp: megabase pairs;
kbp: kilobase pairs.

## Authors’ contribution

I.V.S. and M.A.A. conceived and coordinated the project, and acquired funding for this study. N.A. secured funding for A.Z., J.L. maintained mosquito colonies, performed Hi-C protocol optimization, Hi-C experiments, and Illumina library preparation for *An. arabiensis*. A.S., C.C. and Z.T. did genomic DNA isolation and Oxford Nanopore sequencing. Z.T. performed CQ analysis. Under supervision of M.A.A. and I.V.S., A.Z. and P.A. performed the genome assembly, scaffolding, and validation with the contribution from V.L. N.A. and P.A. supervised A.Z. The manuscript was prepared and approved by all the authors.

## Acknowledgments

The following reagents were obtained through the Biodefense and Emerging Infections Research Resources Repository, NIAID, NIH: *An. arabiensis*, Strain DONGOLA 2Ra, 2Rb/b and 3R, MRA-1235, contributed by Ellen M. Dotson; *An. coluzzii*, Strain MOPTI, Eggs, MRA-763, contributed by Gregory C. Lanzaro. The authors thank Adam Kai Leung Wong for his invaluable help in installing and troubleshooting various software packages on the Colonial One high-performance computing cluster at the George Washington University. We thank Janet Webster and Kristi DeCourcy for editing the manuscript.

