## Additional file 2 for "Chromosome-level genome assemblies of the malaria vectors *Anopheles coluzzii* and *Anopheles arabiensis*"

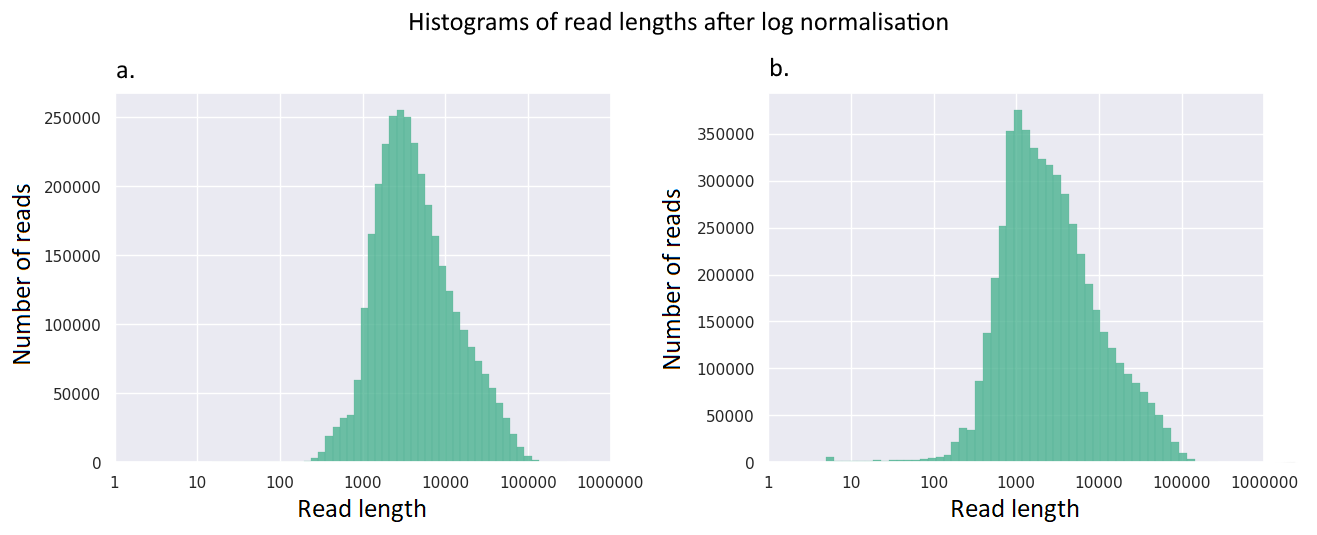
**Additional file 2.** The histogram of read length after log normalization for Nanopore reads from **(a)** *An. coluzzii* and **(b)** *An. arabiensis*.
