## Additional file 3 for "Chromosome-level genome assemblies of the malaria vectors *Anopheles coluzzii* and *Anopheles arabiensis*"

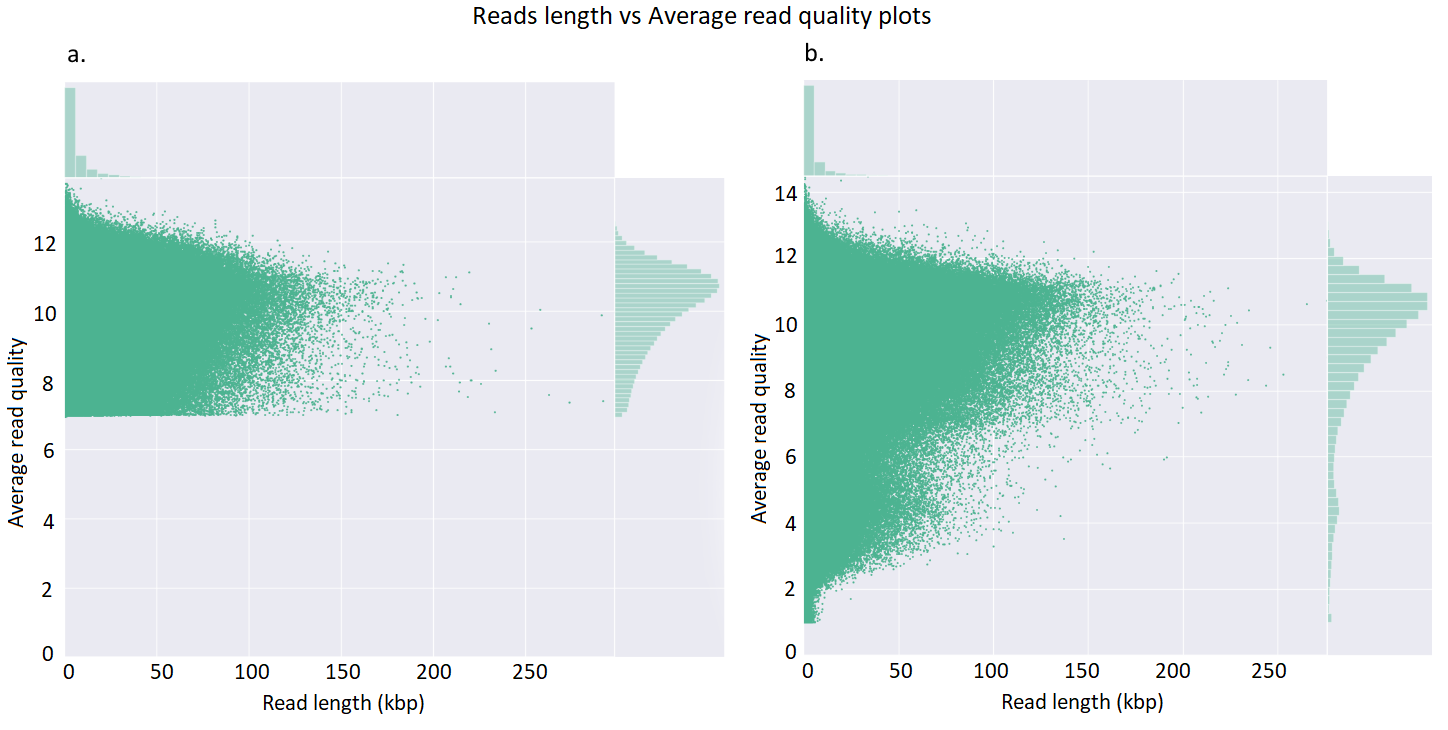

**Additional file 3.** The plot of average read quality for Nanopore reads obtained from **(a)** *An. coluzzii* and **(b)** *An. arabiensis*.
