## Additional file 4 for "Chromosome-level genome assemblies of the malaria vectors *Anopheles coluzzii* and *Anopheles arabiensis*"

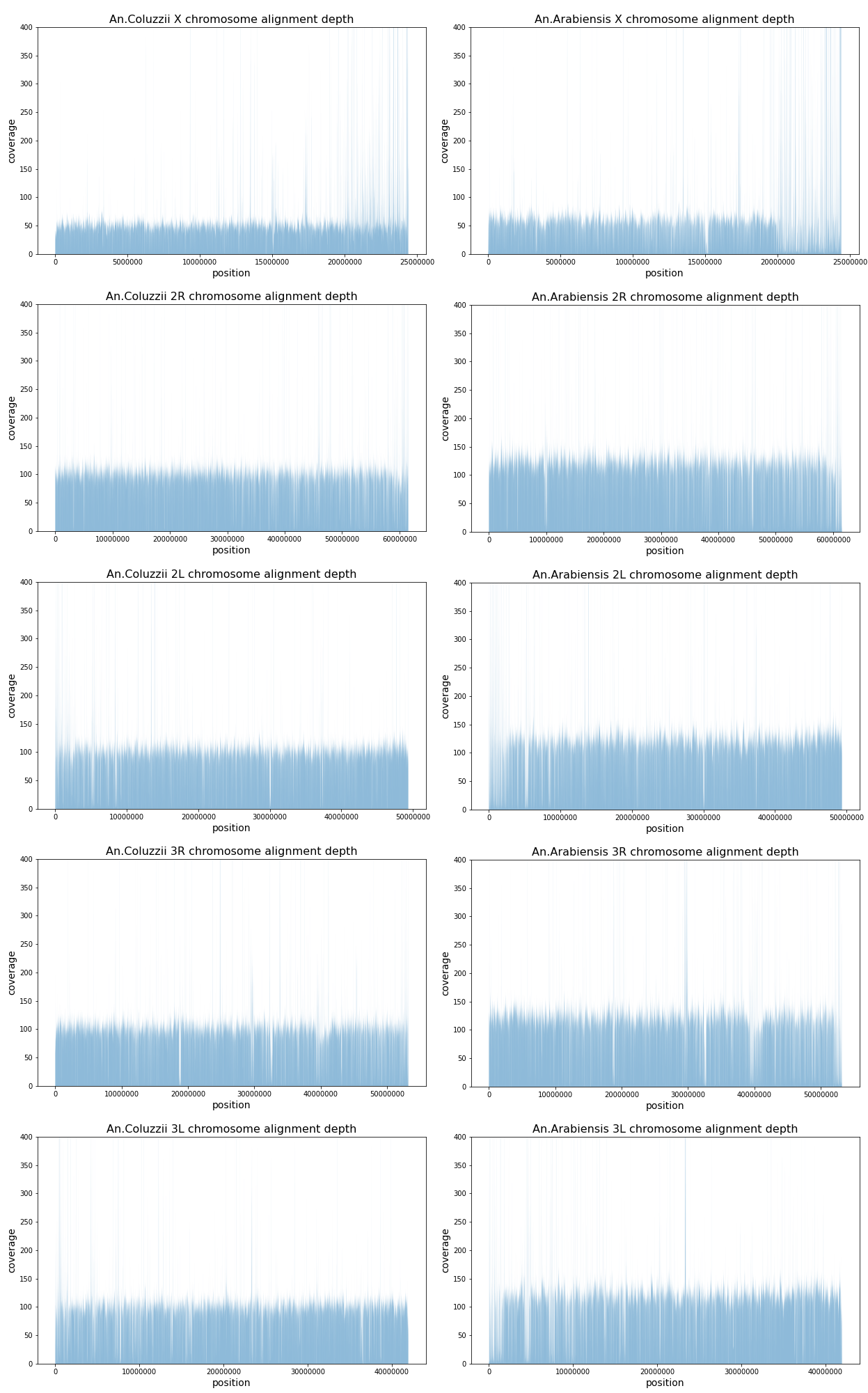

**Additional file 4.** The alignment depth of Nanopore reads from *An. coluzzii* (left column) and *An. arabiensis* (right column) to the *An. gambiae* (AgamP4) genome.
