## Additional file 5 for "Chromosome-level genome assemblies of the malaria vectors *Anopheles coluzzii* and *Anopheles arabiensis*"

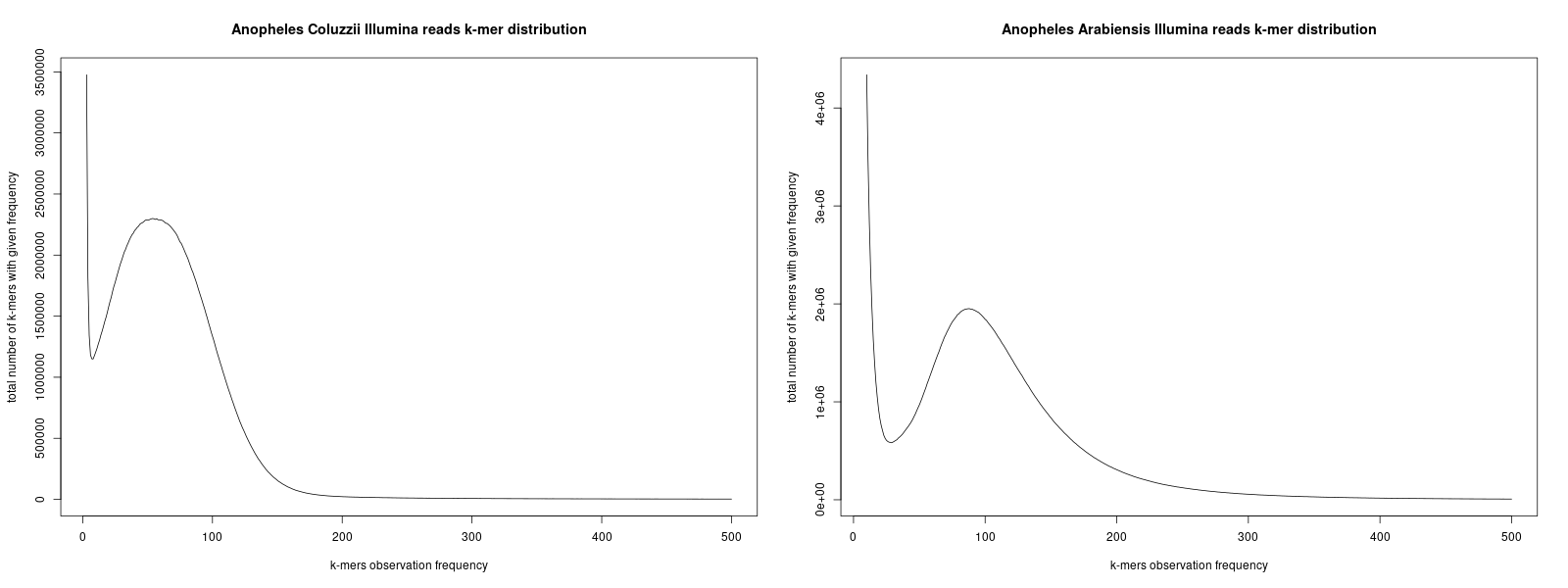

**Additional file 5.** Distribution of 19-mers for *An. coluzzii* (left panel) and *An. arabiensis* (right panel) computed by Jellyfish.
