## Additional file 12 for "Chromosome-level genome assemblies of the malaria vectors *Anopheles coluzzii* and *Anopheles arabiensis*"

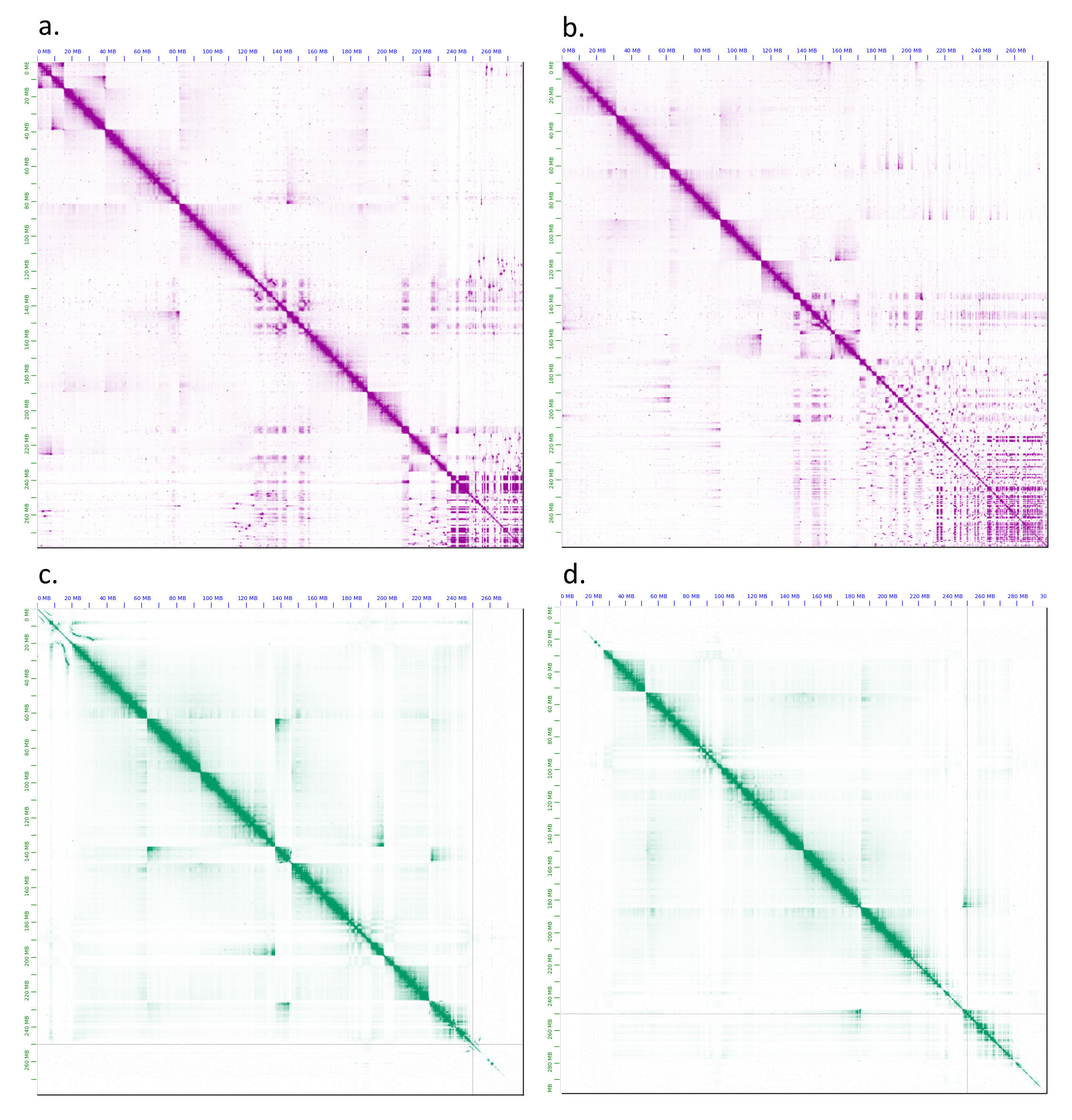

**Additional file 12.** The initial heat maps of Hi-C contact information for the *An. arabiensis* genome assemblies obtained by **(a)** SALSA 2 from the Canu contig assembly, **(b)** SALSA 2 from the Canu unitig assembly, **(c)** 3D-DNA from the Canu contig assembly, and **(d)** 3D-DNA from the Canu unitig assembly. The heat maps are produced by JBAT.
