## Additional file 13 for "Chromosome-level genome assemblies of the malaria vectors *Anopheles coluzzii* and *Anopheles arabiensis*"

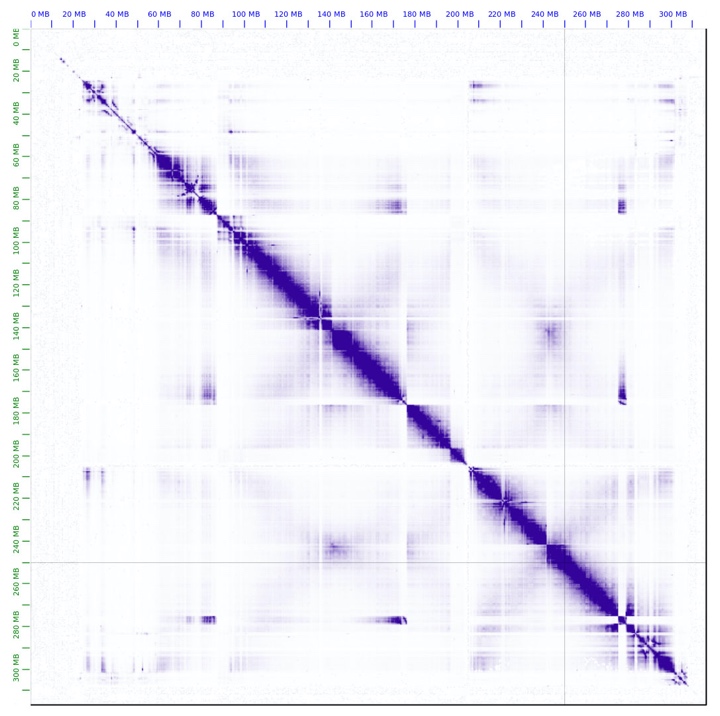

**Additional file 13.** The Hi-C contact heat map for the 3D-DNA scaffolds of the *An. coluzzii* assembly before manual correction. The heat map is produced by JBAT.
