## Additional file 14 for "Chromosome-level genome assemblies of the malaria vectors *Anopheles coluzzii* and *Anopheles arabiensis*"

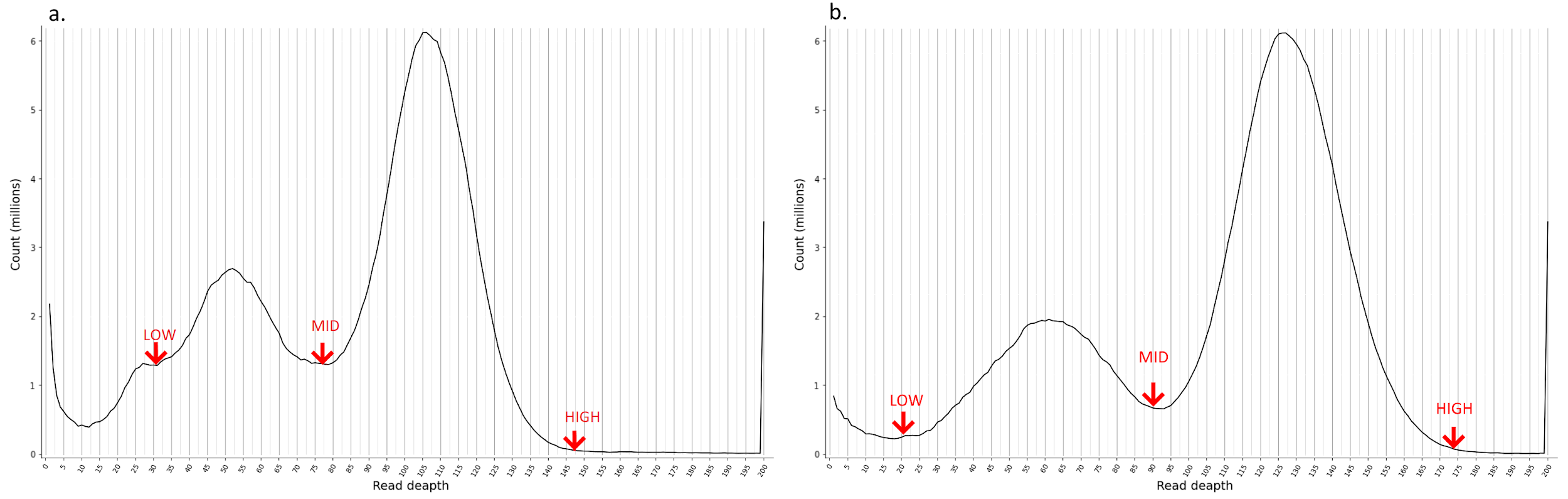

**Additional file 14.** The read depth histogram obtained by Purge Haplo for the *An. coluzzii* **(a)** and *An. arabiensis* **(b)** assemblies. The cut-offs were manually selected (red arrows in the histograms): 30, 78, and 132 for *An. сoluzzii* and 25, 93, and 160 for *An. arabiensis*.
