## Additional file 16 for "Chromosome-level genome assemblies of the malaria vectors *Anopheles coluzzii* and *Anopheles arabiensis*"

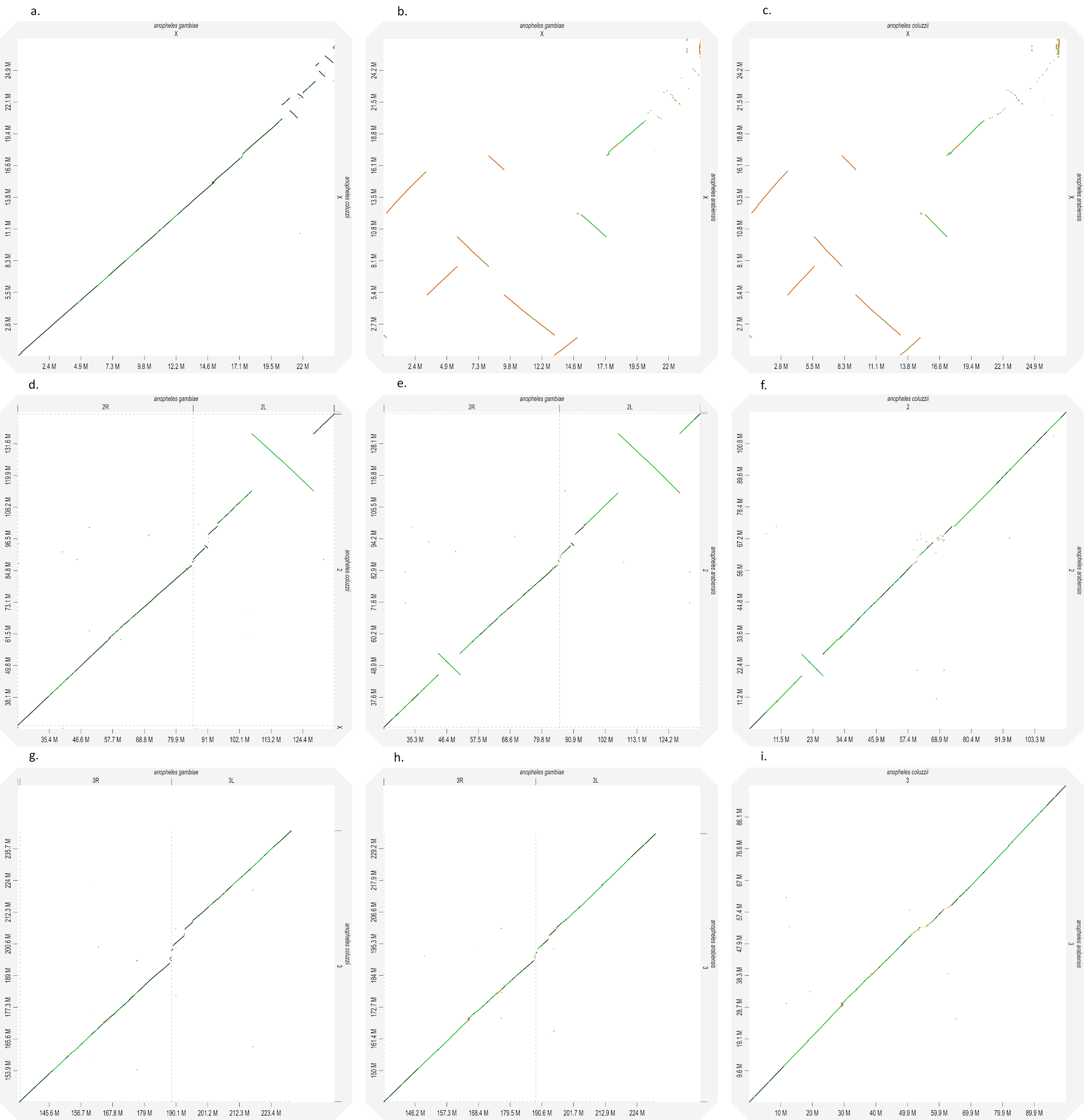

**Additional file 16.** Whole-genome pairwise alignment dot-plots (produced by D-Genies v1.2.0) between the scaffolds corresponding to the chromosomes X **(a, b, c)**, 2 **(d, e, f)**, 3 **(g, h, i)**. Alignments between the *An. gambiae* and *An. coluzzii* scaffolds*,* the *An. gambiae* and *An. arabiensis* scaffolds, the *An. coluzzii* and *An. arabiensis* scaffolds are show.
