## Additional file 20 for "Chromosome-level genome assemblies of the malaria vectors *Anopheles coluzzii* and *Anopheles arabiensis*"

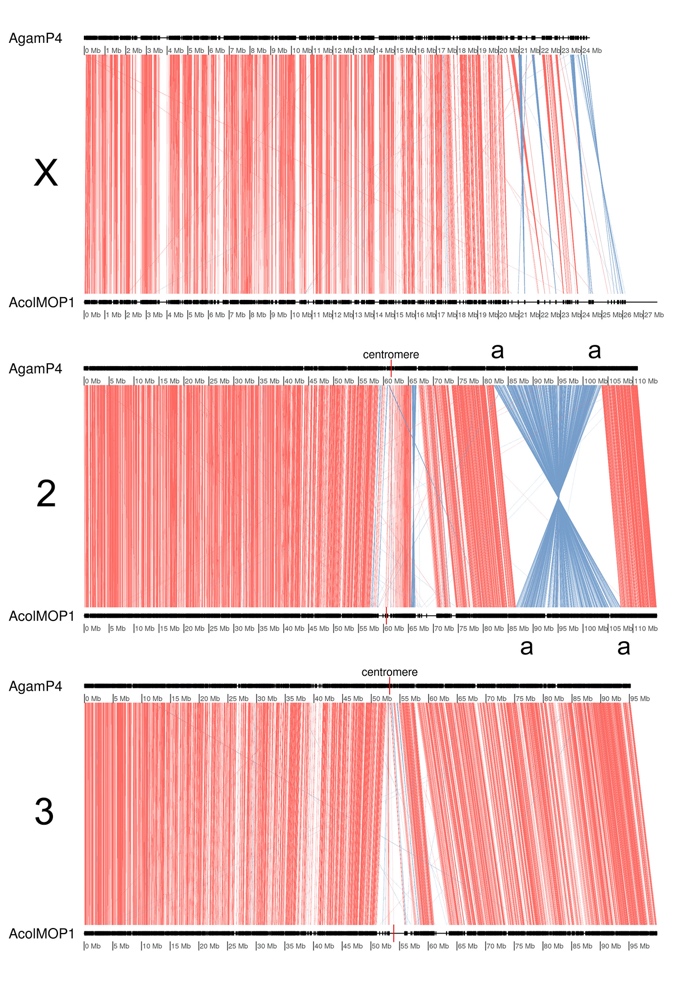

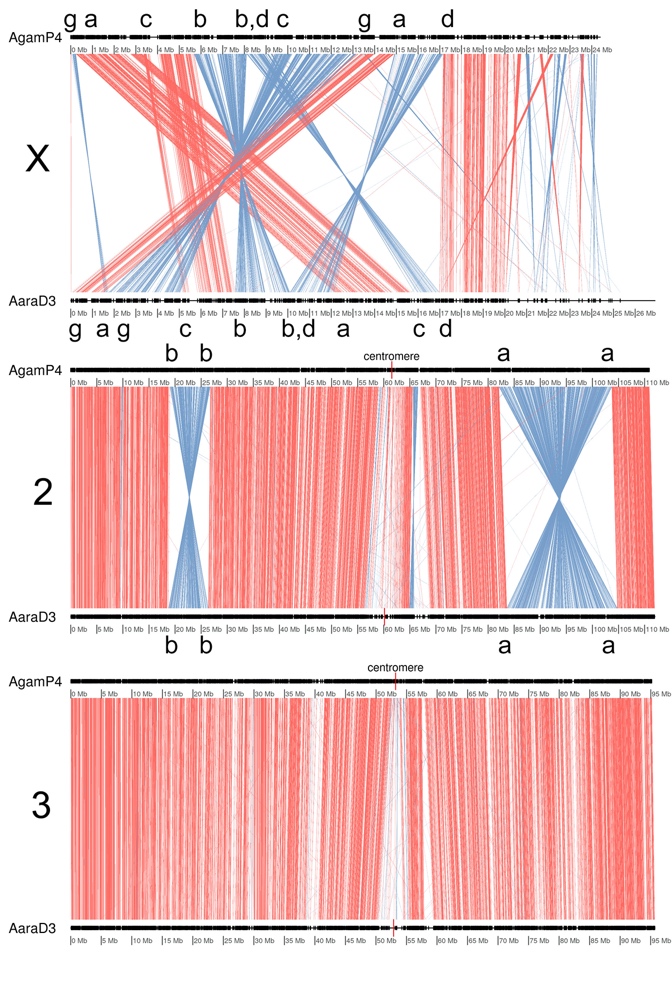

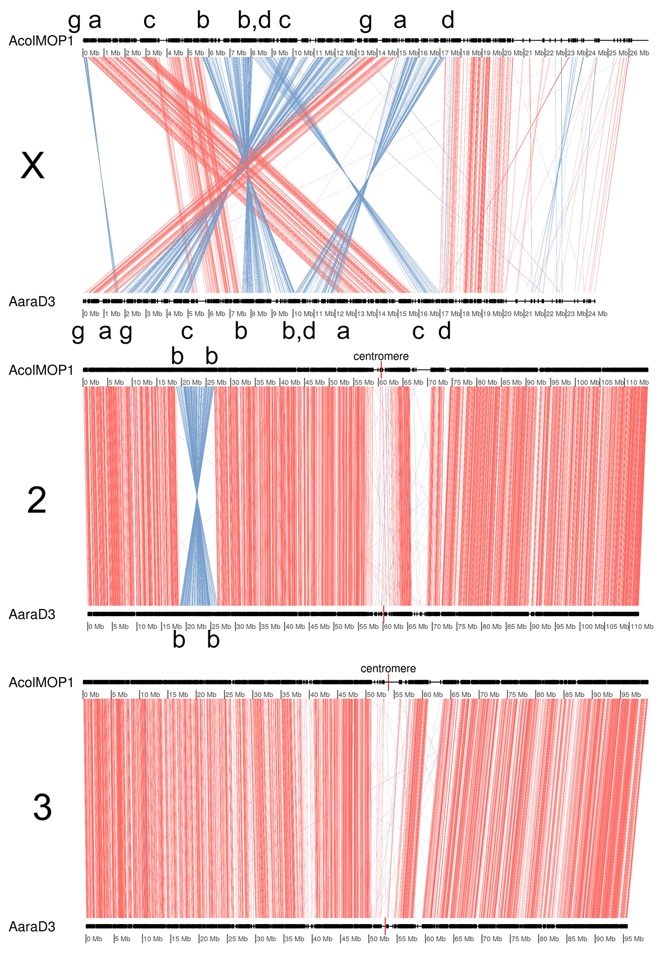

**Additional file 20.** Whole-genome pairwise alignments produced by genoPlotR between chromosomes of *An. arabiensis, An. coluzzii, and An. gambiae* Left panel: AgamP4 and AcolMOP1. Middle panel: AgamP4 and AaraD3. Right panel: AcolMOP1 and AaraD3.

The inversion breakpoints are shown with small letters.
