## Additional file 21 for "Chromosome-level genome assemblies of the malaria vectors *Anopheles coluzzii* and *Anopheles arabiensis*"

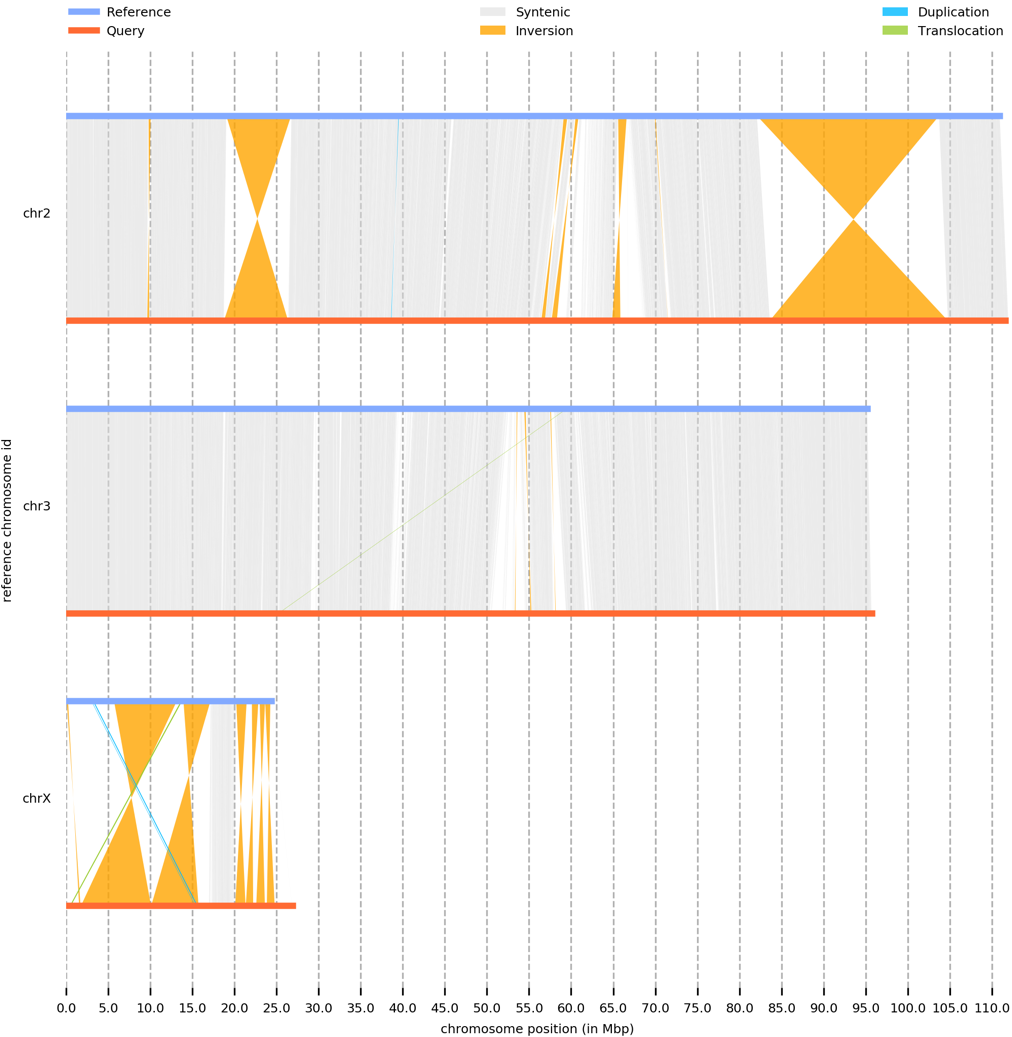

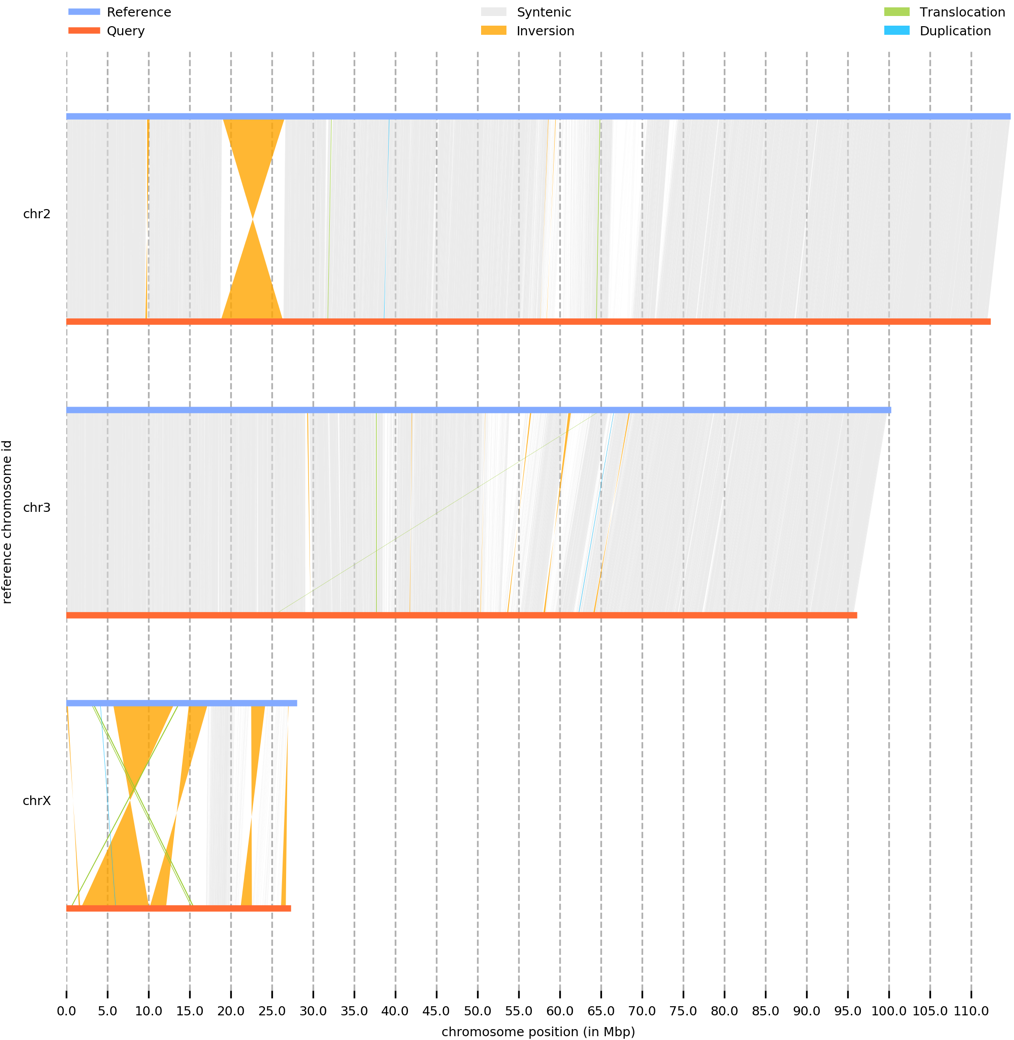

**Additional file 21.** Whole-genome pairwise alignments produced by SyRi of *An. arabiensis* chromosomes (query) to *An. gambiae* (reference)*,* and *An. coluzzii* (reference) chromosomes. Left panel: AgamP4 and AaraD3. Right panel: AcolMOP1 and AaraD3.
