## Additional file 24 for "Chromosome-level genome assemblies of the malaria vectors *Anopheles coluzzii* and *Anopheles arabiensis*"

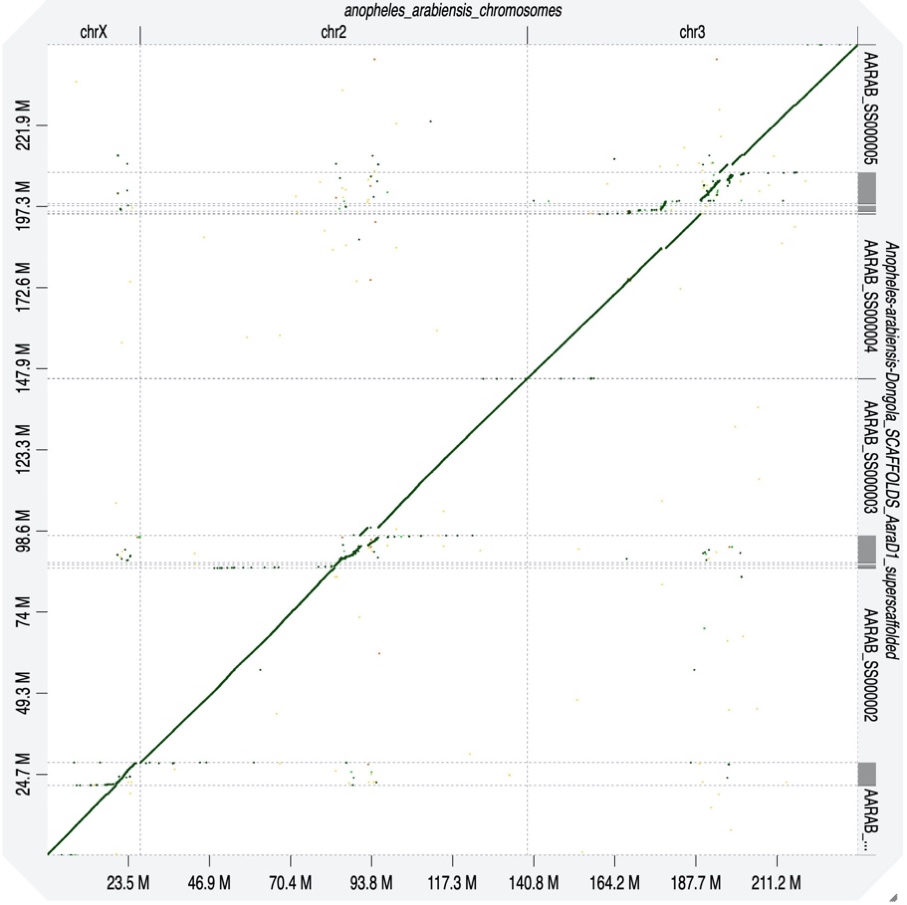

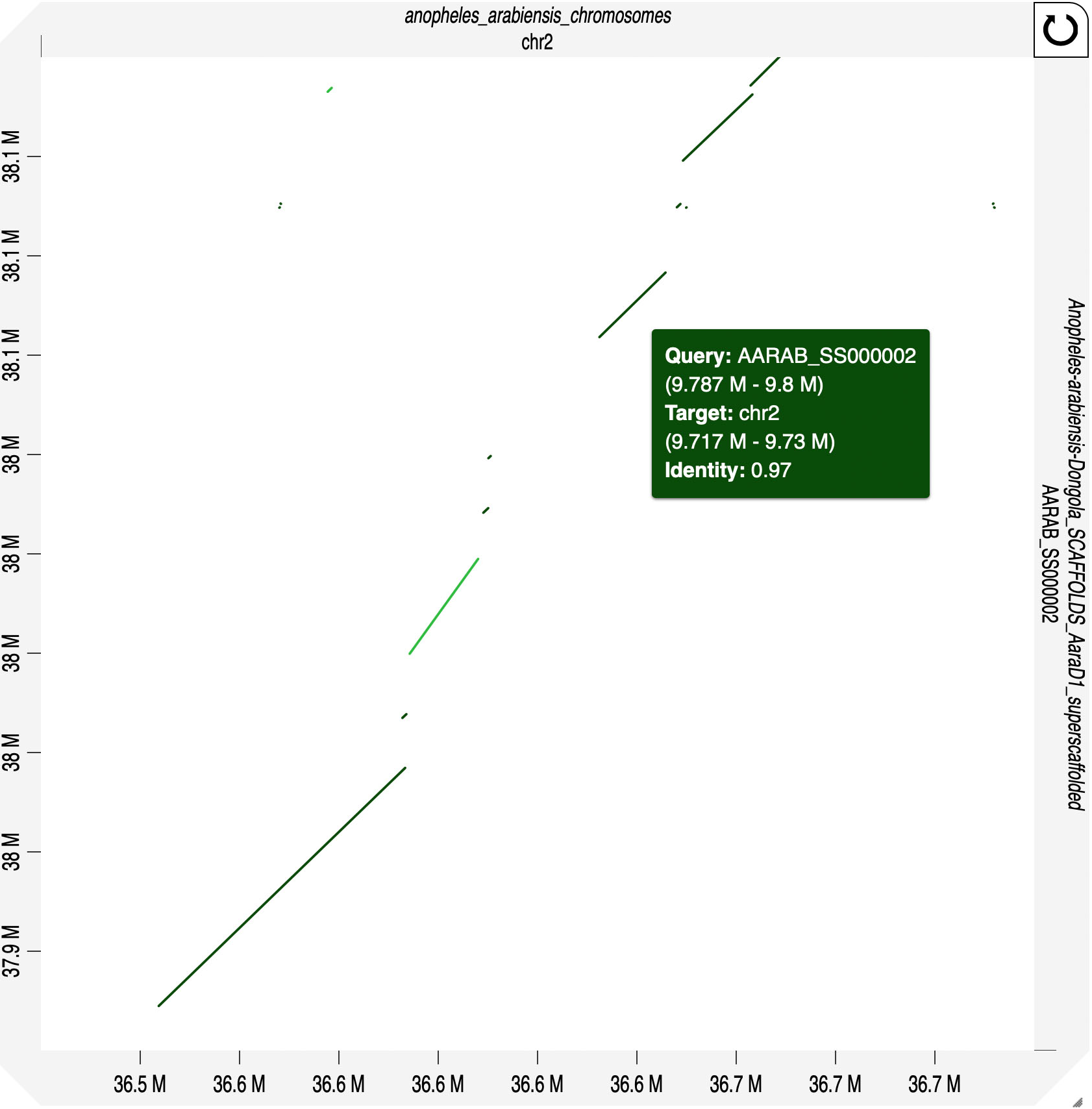

**Additional file 24.** Pairwise dot-plot alignment between the AaraD3 and AaraD2 (AaraD1 superscaffolded) assemblies produced by D-Genies v1.2.0. Top panel: Whole-genome pairwise alignment. Bottom panel: Collinearity in the alignment of the region with the new 2R microinversion.
