## Additional file 25 for "Chromosome-level genome assemblies of the malaria vectors *Anopheles coluzzii* and *Anopheles arabiensis*"

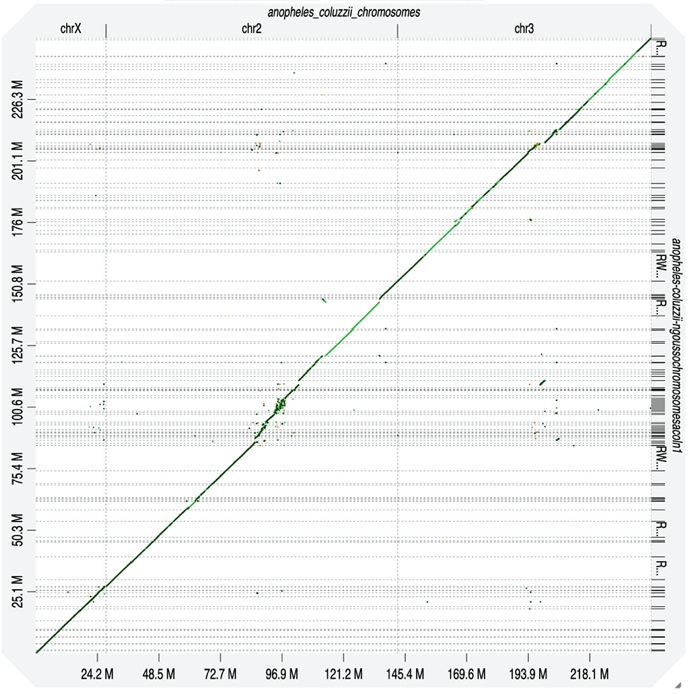

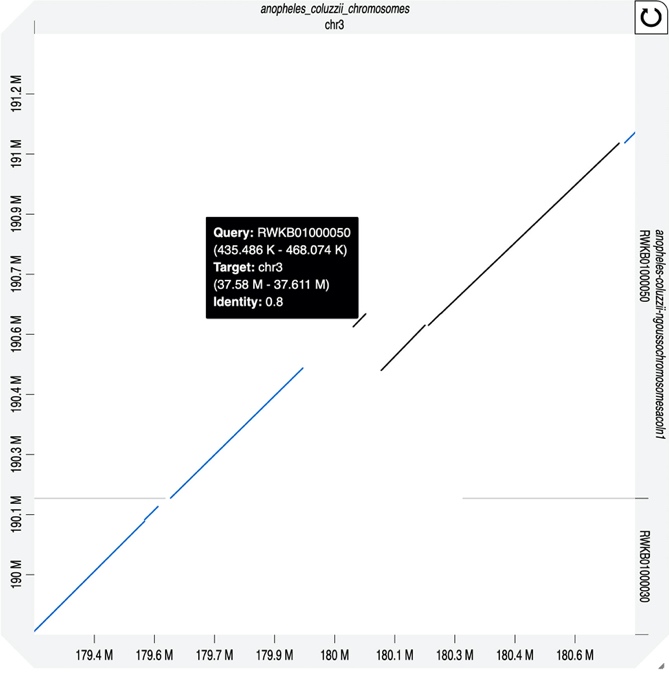

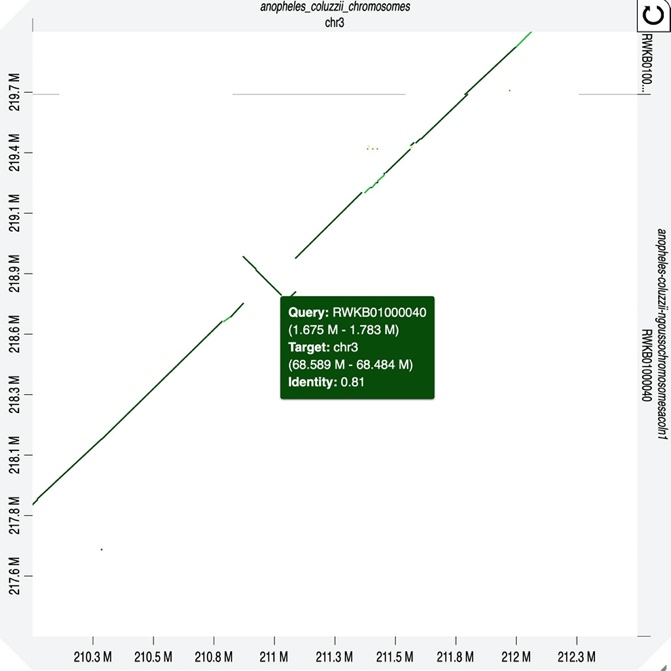

**Additional file 25.** Pairwise dot-plot alignment between the AcolMOP1 and reference-guided scaffolded AcolN1 assemblies produced by D-Genies v1.2.0. Left panel: Whole-genome pairwise alignment. Middle panel: The alignment of the region with the new 3R translocation. Right panel: The alignment of the region with the new 3L microinversion.
