## Additional file 27 for "Chromosome-level genome assemblies of the malaria vectors *Anopheles coluzzii* and *Anopheles arabiensis*"

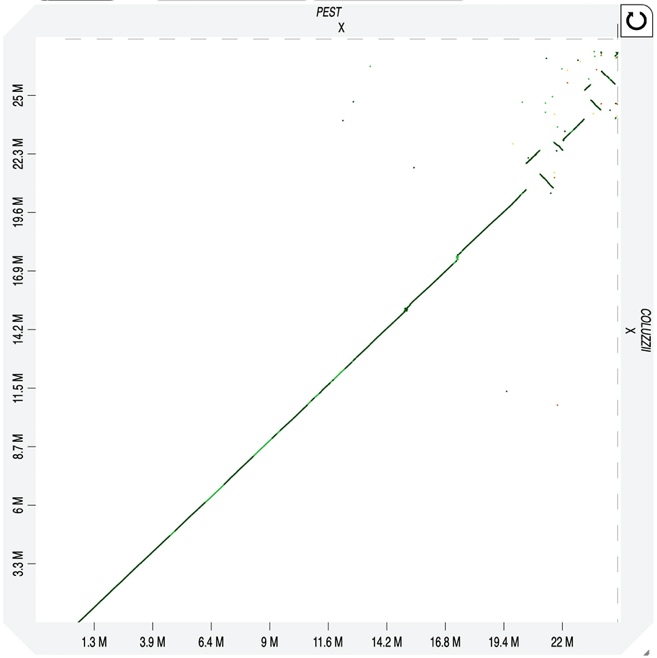

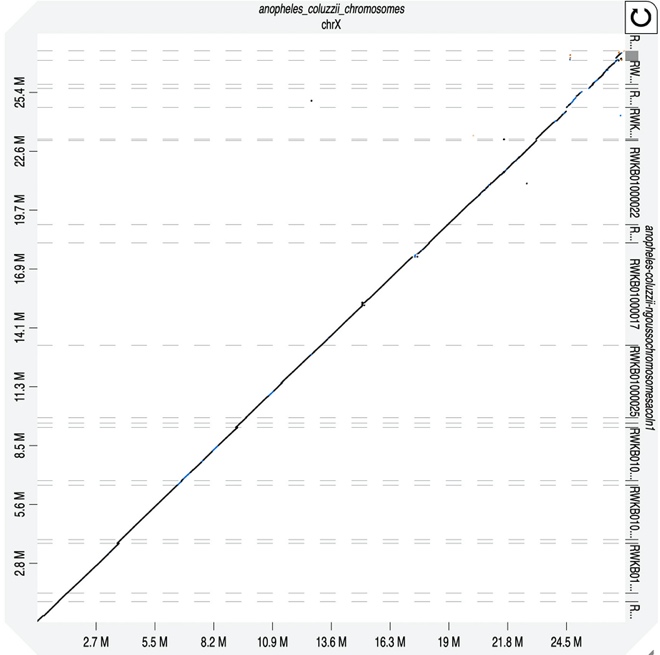

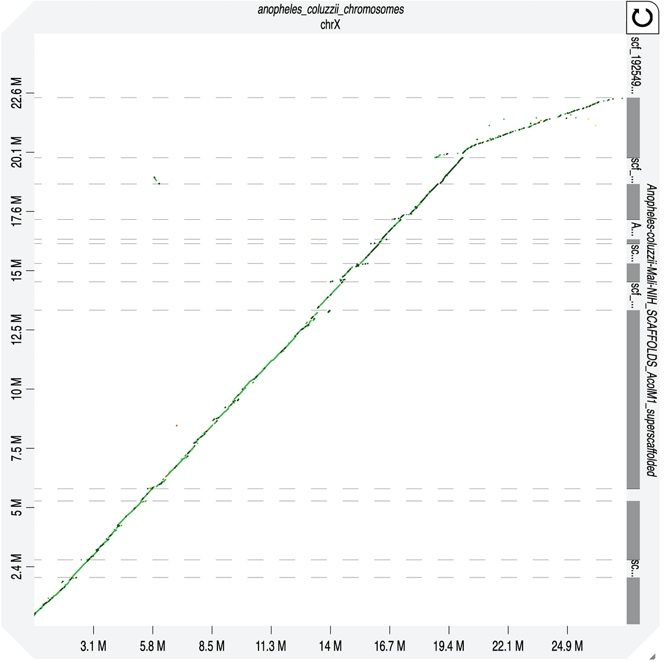

**Additional file 27.** Pairwise dot-plot alignment between the X chromosomes produced by D-Genies v1.2.0. Left panel: AcolMOP1 and AgamP4. Middle panel: AcolMOP1 and AcolN1. Right panel: AcolMOP1 and AcolM2.
